# Early structural connectivity within the sensorimotor network: deviations related to prematurity and association to neurodevelopmental outcome

**DOI:** 10.1101/2022.05.04.490626

**Authors:** Neumane Sara, Gondova Andrea, Leprince Yann, Hertz-Pannier Lucie, Arichi Tomoki, Dubois Jessica

**Author notes:** These authors have contributed equally to this work and share first authorship (Neumane Sara, Gondova Andrea).

## Abstract

The sensorimotor (SM) network is crucial for optimal neurodevelopment. However, undergoing rapid maturation during the perinatal period, it is particularly vulnerable to preterm birth.

Our work explores the prematurity impact on the microstructure and maturation of primary SM white matter (WM) tracts at term-equivalent age (TEA) and evaluates the relationships between these alterations and neurodevelopmental outcome.

We analyzed diffusion MRI data from the developing Human Connectome Project (dHCP) database: 59 preterm (PT) low-risk infants scanned near TEA, compared to a control group of full-term (FT) neonates paired for age at MRI and sex. We dissected pairwise connections between primary SM cortices and subcortical structures using probabilistic tractography and evaluated their microstructure with diffusion tensor imaging (DTI) and neurite orientation dispersion and density imaging (NODDI) models. In addition to tract-specific univariate analyses of diffusion metrics, we computed a *maturational distance related to prematurity* based on a multi-parametric Mahalanobis distance of each PT infant relative to the FT group. Finally, we evaluated the relationships between this distance and Bayley Scales of Infant and Toddler Development (BSID-III) scaled scores at 18 months corrected age.

Our results confirm important microstructural differences in SM tracts between PT and FT infants, with effects increasing with lower gestational age at birth. Additionally, comparisons of maturational distances highlight that prematurity has a differential effect on SM tracts which follows the established WM caudo-rostral developmental pattern. Our results suggest a particular vulnerability of projections involving the primary sensorimotor cortices (S1) and of the most rostral tracts, with cortico-cortical and S1-Lenticular tracts presenting the highest alterations at TEA. Finally, NODDI-derived maturational distances of specific tracts seem related to fine motor and cognitive scores.

This study expands the understanding of the impact of early WM alterations in the emerging SM network on long-term neurodevelopment. In the future, related approaches have potential to lead to the development of neuroimaging markers for neurodevelopmental disorders, with special interest for subtle neuromotor impairments frequently observed in preterm-born children.

## Introduction

The cerebral somatosensory and motor systems consist of distributed networks of specialized interconnected cortical and subcortical gray matter (GM) regions, interacting through white matter (WM) tracts, that support a wide variety of sensory and motor functions that are essential to nearly every human behavior across the lifespan. In somatosensation, inputs from peripheral receptors are first conveyed by peripheral nerves, then through the spinal cord to the *brainstem* dorsal column nuclei. These nuclei further connect to the thalamus which sends projections to cortical somatosensory areas, particularly the *primary somatosensory cortex* (S1) located on the *postcentral gyrus*. On the other hand, the *primary motor cortex* (M1) in the *precentral gyrus*, is critical for motor behavior, exerting its influence over the body’ s muscles through its output to a variety of descending pathways, the main being the direct cortical innervation of motoneurons via the corticospinal tract (CST). S1 and M1 cortices are reciprocally connected, *directly* via short-range intra-hemispheric and homotopic interhemispheric pathways, and *indirectly* via some cortico-subcortical pathways predominately involving the *thalamus* and the *basal ganglia* (BG). The BG notably include the *caudate nucleus, putamen* and globus pallidus: the first two functionally constitute the striatum (receiving most of the BG inputs), while the last two are grouped anatomically in the lenticular nucleus, with the globus pallidus representing one of the key output structures of the BG (Leisman et al., 2014).

Interactions between somatosensory and motor systems, observable in mature brains (Hatsopoulos and Suminski, 2011; Tomasino and Gremese, 2016), are particularly important during the early stages of neurodevelopment. The late second and third trimesters of gestation, as well as the neonatal period, are a critical phase for the dynamic refinement and complex maturation of brain networks through several complex processes (Dubois et al., 2014; Kostović et al., 2019), which lay the foundations for later neurodevelopment (Gilmore et al., 2018). As projection and interhemispheric tracts show rapid growth before 28 weeks of gestational age (wGA) (Keunen et al., 2017), the overall architecture of the *sensorimotor* (SM) network is already established during the preterm period, and is one of the earliest systems starting to mature (Dubois et al., 2014; Ouyang et al., 2019; Machado□Rivas et al., 2021).

During infancy and early childhood, the maturation of brain networks occurs with a spatially heterogeneous and temporally asynchronous progression, generally obeying a *primary-to-higher-order sequence*. This is observed both at the GM and WM level, progressing across 1) primary areas (sensorimotor, visual, auditory systems); 2) adjacent unimodal associative structures; and 3) higher-order associative (multimodal) regions (Flechsig, 1920; Yakovlev and Lecours, 1967; Dubois et al., 2014; Deoni et al., 2015; Kulikova et al., 2015; Croteau-Chonka et al., 2016; Remer et al., 2017; Lebenberg et al., 2019; Yu et al., 2020). The integrity of primary systems, particularly the primary SM network, may therefore play a pivotal role for the optimal development of secondary and associative networks in their earliest stages and for organizing the structural and functional connectome throughout the neonatal period (Ball et al., 2014; van den Heuvel et al., 2015; Zhao et al., 2019).

In this context, the preterm period and first postnatal months are crucial for SM system maturation, as emphasized by the adverse effects of preterm birth (before 37wGA) on neurodevelopment. Here, the sudden need to adapt to extra-uterine life during a period of intense maturation for the primary brain networks, likely does not provide the optimal conditions for physiological neurodevelopmental mechanisms, resulting in (sometimes subtle) structural and/or functional abnormalities (Suzuki, 2007). The related diffuse cerebral dysmaturation disorder (Back, 2015; Volpe, 2021) interferes with whole-brain developmental processes and alters the integrity of the emerging neural networks (Suzuki, 2007; Back, 2015), with early maturing regions suffering the greatest adverse effects with increased prematurity (lower GA at birth) (Knight et al., 2018).

Magnetic resonance imaging (MRI) including diffusion MRI has been extensively used to evaluate the consequences of preterm birth on brain development. Even in the absence of focal cerebral lesions, prematurity is associated with disturbances in brain growth, in particular in GM structures including the BG and thalamus (Keunen et al., 2012; Padilla et al., 2015; Loh et al., 2020), and pervasive widespread abnormalities in GM and WM microstructure, maturation and connectivity (Ball et al., 2013b; Batalle et al., 2018). In particular, the WM of preterm infants at term-equivalent age (TEA) has a more “immature” microstructural profile compared with term-born neonates, consistent with delayed and/or disrupted WM development and maturation (Thompson et al., 2011; Kelly et al., 2016a, 2020).

Preterm birth is associated with an increased risk of diverse neurodevelopmental impairments (Johnson and Marlow, 2017; Twilhaar et al., 2018; Pierrat et al., 2021), and the extent of early WM abnormalities (even in the absence of overt brain lesion) has been related to poorer neurodevelopmental outcome (Duerden et al., 2015; Barnett et al., 2018; Kelly et al., 2020; Pannek et al., 2020). Moreover, preterm infants present a higher risk for impaired neuromotor function (Williams et al., 2010; Odd et al., 2013; Spittle and Orton, 2014), that can manifest as early alterations in spontaneous movements which are visible from the first post-natal weeks (Peyton et al., 2020), and later, with poorer fine and gross motor skills compared with term-born controls (Evensen et al., 2020). The relationships between SM network structural alterations and neuromotor impairment in *low-risk preterm* infants (including moderate to late preterm and/or preterm babies without perinatal brain injury) have been less systematically explored, although long-lasting WM alterations in SM tracts have been observed during childhood and adolescence in these populations (Groeschel et al., 2014; Dewey et al., 2019; Thompson et al., 2020).

In this context, further studies in these infants, focusing on the SM network, are required to better explore the WM dysmaturation associated with the common SM and global neurodevelopmental impairments. Specifically, assessing early WM *maturational delays* across the different SM tracts and analyzing the correlation with further outcome would enable a better understanding of the pathophysiology of the disorders resulting from deviations in typical developmental processes.

With these factors in mind, we aimed to study how preterm birth impacts SM network maturation at TEA even in the absence of an overt lesion, and more precisely, to analyze the likely differential effect of the GA at birth on WM microstructural characteristics across the primary SM tracts, and the potential functional consequences reflected by later altered outcome. We hypothesized that the SM tracts would show a significant maturation delay in preterm infants compared to full-term neonates, with distinct patterns of altered maturation across the connections between SM structures (based on the assumption that early peripheral stimuli and subcortical structures play a key role in SM network maturation); and that these alterations are associated with motor and global outcomes.

For this purpose, we studied a large cohort of low-risk preterm infants at TEA and full-term neonates from the *developing Human Connectome Project* (dHCP), by investigating WM microstructure and maturation using complementary approaches based on diffusion MRI data.

Diffusion MRI enables study of WM organization during early brain development through detailed estimation of WM bundles with tractography methods and the assessment of WM microstructure with quantitative metrics (Dubois et al., 2014, 2016; Ouyang et al., 2019). In this study, we were interested in two complementary models: Diffusion Tensor Imaging (DTI) and Neurite Orientation Dispersion and Density Imaging (NODDI).

DTI is the most popular but simple model for estimating the degree and direction of water molecule diffusion within each voxel. DTI-derived metric changes have been thought to be related to WM microstructural development and maturational processes (e.g., water content, axonal density, axonal caliber and integrity, premyelination, myelination) (Dubois et al., 2014; Pecheva et al., 2018; Ouyang et al., 2019). Despite the valuable information that DTI provides about the organization and maturation of WM tracts, the interpretation of changes at the microstructural scale remains challenging as DTI metrics are nonspecific and reflect many underlying properties of brain tissue.

The NODDI model (Zhang et al., 2012) aims to evaluate the directional distribution of neurites (axons and dendrites) in a voxel, through a modelling diffusion within 3 tissue compartments (intra-neurite, extra-neurite and cerebro-spinal fluid (CSF) space). This allows the estimation of the Neurite Density Index (NDI), which quantifies the packing density of axons or dendrites; and the Orientation Dispersion Index (ODI), which assesses the degree of dispersion of neurites resulting from fanning, crossing, etc. (Zhang et al., 2012). NODDI has been previously used to characterize WM characteristics during early development. Globally, NDI increases with maturation in developing WM, and ODI tends to decrease, although this pattern is more complex during the neonatal period (Kunz et al., 2014; Batalle et al., 2017; Kimpton et al., 2021).

Although univariate dMRI approaches, based on individual derived metrics, can reveal developmental changes in the neonatal brain (Kunz et al., 2014; Ouyang et al., 2019), they can hardly fully reflect the complexity of processes involved in early brain maturation (Kostović et al., 2019). Also, quantifying the maturational degree across regions therefore requires comparison of an infant’ s data with a mature reference to account for “intrinsic” microstructural differences (Dubois et al., 2014). As shown in neonatal and pediatric data (Kulikova et al., 2015; Dean et al., 2017; Li et al., 2022), the use of the *multivariate Mahalanobis distance* approach allow to overcome these difficulties, by computing the distance between a subject and a group of reference subjects, with the advantage of taking into account the inter-subject variability in the reference group, as well as correlations between input parameters. Moreover, using Mahalonobis distance with DTI and NODDI complementary metrics might allow to account for the effect of maturation and complex underlying WM microstructural trajectories in the early developmental period and in the context of prematurity.

In this study, we aimed to improve our understanding of the impact of prematurity on the early development of SM tracts, by dissecting an unprecedented set of SM cortico-cortical and cortico-subcortical tracts, and then analyzing the *maturational distances* of *low-risk* preterm infants at TEA compared to full-term reference neonates, using multiparametric analyses and Mahalonobis distance combining different sets of DTI and NODDI metrics. Finally, we aimed to identify the relationship between the aforementioned factors with neurodevelopmental outcome at 18 months of corrected age (mCA).

## Methods

### Subjects

This study included a sample of preterm and full-term neonates taken from the dHCP cohort, collected in London, UK from 2015 to 2020 (http://www.developingconnectome.org/). This project has received UK NHS research ethics committee approval (14/LO/1169, IRAS 138070), and written informed consent was obtained from the parents of all participant infants. From the overall cohort, we identified 59 preterm (PT) infants (33 males, gestational age at birth – GA at birth: median 31.7 weeks, range [23.7w – 36.0w]) scanned near TEA (median post-menstrual age –PMA: 41.3w, range [38.4w – 44.9w]), and a control group of full-term born (FT) infants (GA at birth: median 40.1w, range [37.4w – 42.3w]) matched to the preterm population on age at MRI and sex. Preterm infants were subdivided into infants born extremely to very preterm (GA at birth <32w, N=33; PT_EV_ group) or moderate to late preterm (GA at birth ≥32w, N=26; PT_ML_ group). The corresponding controls are subsequently noted FT_EVCt_ and FT_MLCt_ respectively. All the included infants were deemed healthy, i.e. were without major brain focal lesions or any overt abnormality of clinical significance on structural MRI as evaluated by an expert pediatric neuroradiologist (dHCP radiological score in the range 1*–*3: https://biomedia.github.io/dHCP-release-notes/download.html).

#### Neonatal characteristics at birth

*Obstetric factors* (i.e., multiple pregnancy status, intrauterine growth restriction –IUGR, maternal antenatal steroids and magnesium therapy, delivery method) as well as *infant* additional *characteristics at birth* (i.e., Apgar scores at 1 and 5 min, birth weight, length, and head circumference) were extracted from the dHCP records.

Additionally, *postnatal risk factors* previously recognized to be related with neonatal brain abnormalities, including diffuse and regional WM microstructural alterations (Pogribna et al., 2013; Brouwer et al., 2017; Barnett et al., 2018; Parikh et al., 2021), were also considered. In particular, some NICU variables (i.e., total duration of ventilatory support and oxygen therapy, and parenteral nutrition) were binarized using thresholds established in previous studies (need of mechanical ventilation beyond 7 days, and of parenteral nutrition longer than 21 days) (Brouwer et al., 2017). Sepsis was considered as any situation where an infant received antibiotics, as there was not enough information to retain only confirmed episodes of postnatal sepsis. Additionally, we derived a *neonatal morbidities* binary factor to summarize the presence of at least one of the following 4 morbidities associated with prematurity (or the absence of all 4): chronic lung disease, necrotizing enterocolitis (NEC), retinopathy of prematurity (ROP) and abnormal cranial ultrasonography (cUS). Of note, detailed neonatal medical records were available only for infants admitted to the neonatal intensive care unit (NICU) after birth: 51 PT (86%) and 2 FT (3%, admitted for sepsis treatment, without further complications). Comparisons between PT and FT groups in terms of the described variables and factors were performed with suitable tests (Wilcoxon rank sum test for ordinal and continuous variables; Fisher’ s exact test for binary factors; Pearson’ s Chi-squared test for non-binary nominal factors) in R (version 4.0.5, 2021.03.31).

#### Outcome assessment and infant characteristics at 18 months

*Family socio-economic status* (SES) was measured using the Index of multiple deprivation (IMD) which is a UK geographically defined composite social risk score comprising data on income, employment, health, education, living environment, and crime calculated at 18mCA assessment from the mother’ s home address at the time of birth.

*Neurodevelopmental outcome* was assessed at St Thomas’ Hospital, London by two experienced assessors (a paediatrician and a chartered psychologist) using the Bayley Scales of Infant and Toddler Development, Third Edition –BSID-III (Bayley, 2006). We only considered assessments performed at around 18 mCA (between 17 and 21m), which was available for 44 (75%) PT infants and 53 (90%) FT infants (median age: 18.3m). Five distinct developmental categories: cognition, receptive and expressive language, and fine and gross motor function were assessed yielding age-standardized respective *scaled scores* (mean 10, SD 3), with higher values indicating better infant development and scores lower than 7 indicating developmental delay in that domain.

Comparisons between PT and the FT groups were performed with t-tests corrected for multiple comparisons using Benjamini–Hochberg False Discovery Rate (FDR) correction across scores. The effect of sub-groups (PT_EV,_ PT_ML_, FT_EVCt_, and FT_MLCt_) on neurodevelopmental outcomes was assessed using one-way ANOVA.

Of note, the 5 BSID-III scaled scores can be summarised into the widely used 3 *composite* cognitive, language, and motor scores (mean 100, SD 15). The results of the entire analyses performed using them can be found in the **Supplementary Materials: *BSID-III composite score results*** section.

### MRI data acquired at TEA

MRI data was acquired using a Philips 3-Tesla Achieva scanner (Philips Medical Systems, Best, The Netherlands). All infants were scanned during natural sleep using a neonatal head coil and imaging system optimized for the dHCP study as previously described (Hughes et al., 2017).

We used anatomical and diffusion MRI data available in its pre-processed state from the dHCP database (third release). The *structural* data was a result of acquisition and reconstruction using optimized protocols (Cordero-Grande et al., 2018) leading to super-resolved T2w images with an isotropic spatial voxel size of 0.5mm. Processing followed a dedicated pipeline for segmentation and cortical surface extraction for T2w images of neonatal brains (Makropoulos et al., 2018), with bias-correction, brain extraction, and segmentation using Draw-EM (Developing brain Region Annotation with Expectation Maximisation) algorithm (Makropoulos et al., 2014). White matter surface (inner cortical surface) meshes provided within the dHCP database were used for the segmentation of cortical regions of interest (ROIs), while volumetric segmentations were directly used to extract subcortical ROIs (cf. section ***Delineation of regions of interest***). Additionally, derived hemispheric, WM and cortical masks (also referred to as cortical ribbons) were also used for the tractography analysis (cf. section ***SM connectivity reconstruction***).

Acquisition and reconstruction of the *diffusion* data followed a multi-shell high angular resolution diffusion imaging (HARDI) protocol with 4 b-shells (b=0□s/mm^2^: 20 repeats; and b= 400, 1000, 2600 s/mm^2^: 64, 88 and 128 directions respectively) (Hutter et al., 2018) and was pre-processed with correction for motion artefacts and slice-to-volume reconstruction using the SHARD approach, leading to an isotropic voxel size of 1.5mm (Christiaens et al., 2021). Pre-processed data was used for the fitting of diffusion models (cf. section ***Estimation of diffusion models***) and for the tractography analysis (cf. section ***SM connectivity reconstruction***).

### Assessment of SM network microstructure

To estimate WM microstructural characteristics within the SM network, we first quantified complementary diffusion metrics from the available diffusion data. We then reconstructed the structural connections between pairs of anatomically defined SM regions, including cortical primary SM cortices and key sub-cortical structures, using probabilistic tractography. The diffusion metrics were then extracted for the selected connections of interest and used to study developmental differences between the cohort subgroups.

#### Estimation of diffusion models

DTI model was fitted to the diffusion data using a single shell (b=1000 s/mm^2^) and calculated with FSL’ s DTIFIT. The choice of using only a single b-value was made because the utility of including more diffusion directions may be outweighed by the non-Gaussian contribution of high b-value acquisitions (Pines et al., 2020). DTI maps were computed for 4 metrics: Fractional Anisotropy (FA) and Mean Diffusivity (MD) which are a composite of Axial Diffusivity (AD) and Radial Diffusivity (RD).

Additionally, multi-shell data were used to derive the NDI and ODI maps from the NODDI model (Zhang et al., 2012) using the CUDA 9.1 Diffusion Modelling Toolbox (cuDIMOT) NODDI Watson model implementation for GPUs (Hernandez-Fernandez et al., 2019). We used the MCMC optimization algorithm and default settings to fit the NODDI model to our infant data. The NODDI-derived maps were then post-processed to reduce the observed noise. Briefly, we used ODI maps to detect possible errors using an alpha- trimming strategy. The voxels presenting values outside the threshold range (fixed upper value of 0.95 and the lower limit being the first groove of the histogram of values) were either i) normalized by the immediate surrounding values (i.e. the mean of the voxel’ s immediate environment after the removal of extreme values), or ii) set to 0, if no voxels in the ‘ normal’ range were found in their environment. The same erroneous voxels were also corrected in NDI maps in the same fashion.

#### Delineation of regions of interest

Pre-processed structural data was used to parcellate 13 ROIs (6 in each hemisphere and 1 bilateral, **Figure 1.A**). *Three cortical* ROIs were defined on the cortical surface of each hemisphere using the M-CRIB-S surface-based parcellation tool optimized for the term-born neonates (Adamson et al., 2020) whose labelling scheme replicates the Desikan-Killiany-Tourville (DKT) atlas (Klein and Tourville, 2012): the postcentral gyrus as the anatomical proxy of the lateral portion of the primary somatosensory cortex (hereafter referred to as *S1* for the sake of simplification), the precentral gyrus as the lateral portion of the primary motor cortex (referred to as *M1*), and the paracentral lobule (referred to as ParaC) corresponding to the medial surface of the hemisphere in the continuation of the precentral and postcentral gyri, including the medial portions of the primary sensori-motor cortices. Individual surface ROIs were then projected to the cortical ribbon defined in the anatomical volumes, and further dilated by one voxel into the WM to ease the tractography process.

**Figure 1.**
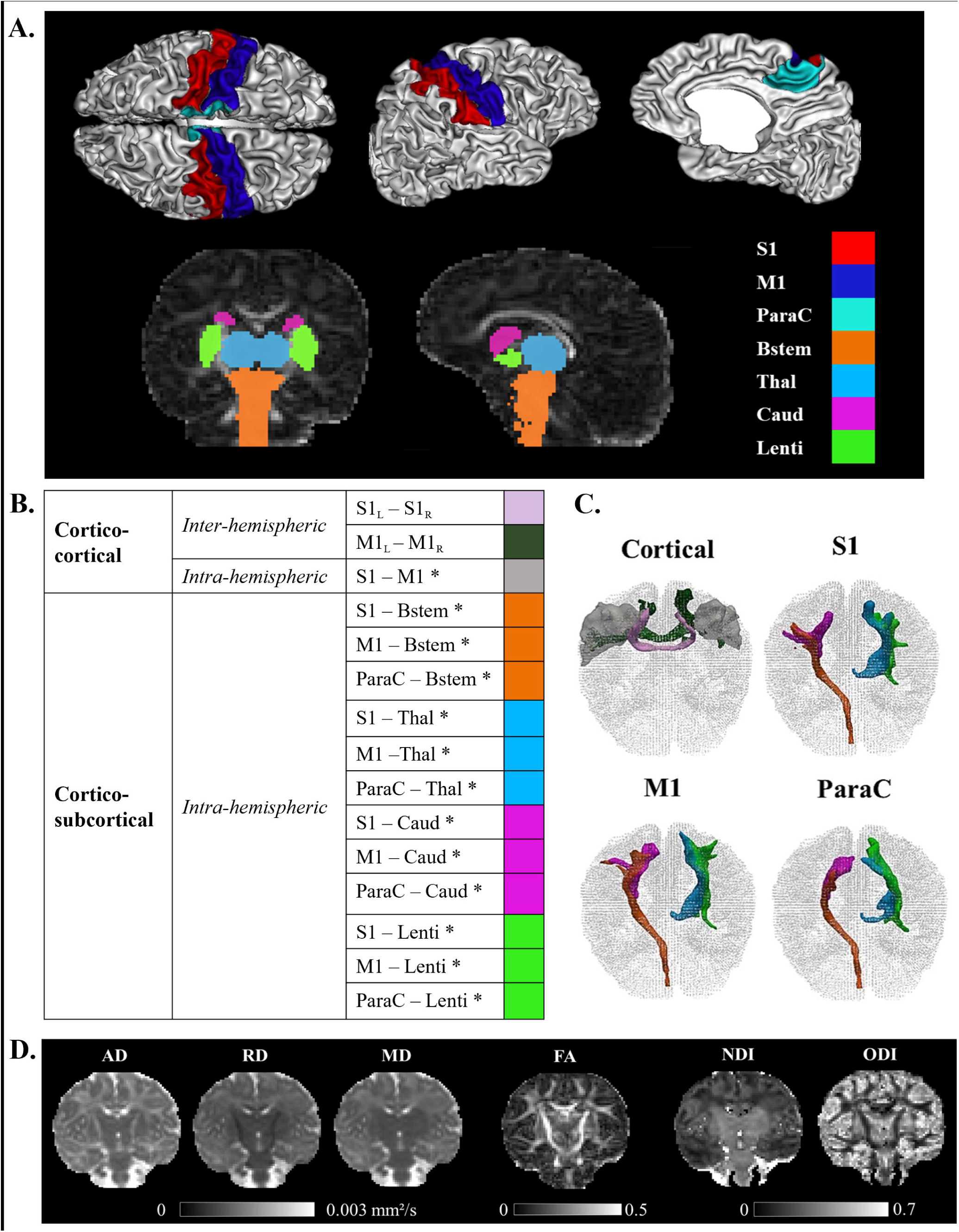
Regions of interest, sensorimotor tracts, and diffusivity metrics for a representative full-term infant (GA at birth 40.4w, PMA at MRI 44.1w). **A**. Visualization of the cortical and subcortical ROIs used as tractography seeds. **B**. List of the SM tracts of interest. **C**. 3D reconstructions of SM tracts. **D**. Metric maps resulting from DTI (AD, RD, MD, FA) and NODDI (NDI, ODI) models. *Legends*. GA: gestational age (in weeks), PMA: post-menstrual age (in weeks). ROIs: regions of interest. SM: sensorimotor. S1: lateral portion of the primary somatosensory cortex (postcentral gyrus); M1: lateral portion of the primary motor cortex (precentral gyrus); ParaC: medial portions of the primary sensorimotor cortices (paracentral area). Bstem: brainstem; Thal: thalamus; Caud: caudate nucleus; Lenti: lentiform nucleus. L: left; R: right. * Intra-hemispheric tracts, evaluated in left and right hemispheres separately. DTI (Diffusion Tensor Imaging) metrics: AD (Axial Diffusivity), RD (Radial Diffusivity), MD (Mean Diffusivity), FA (Fractional Anisotropy); NODDI (Neurite Orientation Dispersion and Density Imaging) metrics: NDI (Neurite Density Index), ODI (Orientation Dispersion Index).

*Subcortical* ROIs were defined using the subcortical grey matter parcellation based on Draw-EM algorithm segmentation (Makropoulos et al., 2014) provided within the dHCP data release, namely medial brainstem (Bstem) and for each hemisphere: thalamus (Thal, fusing high and low intensity regions), caudate nucleus (Caud) and lentiform nucleus (Lenti). These ROIs were used as seeds for the tractography.

Cortical ROIs (bilateral M1, S1, and ParaC regions) and subcortical ROIs (brainstem and bilateral thalamus, caudate and lenticular nuclei) tractography seeds were aligned to the diffusion space with FSL 6.0’ s FLIRT.

#### SM connectivity reconstruction

For each subject, probabilistic tractography estimating multiple diffusion orientations within a voxel (Behrens et al., 2007) was used to reconstruct connections between the selected ROI pairs (designated as *tracts* thereafter). Briefly, for each subject, we first modelled crossing fibers within each voxel of the multi-shell diffusion data using a GPU accelerated version of FSL’ s Bayesian Estimation of Diffusion Parameters Obtained using Sampling Techniques modelling Crossing Fibres (BEDPOSTX), with default settings apart from the deconvolution model with zeppelins (Hernández et al., 2013). Then, the pre-selected ROIs were used as seed masks to derive region-to-region structural connections using the GPU implementation of the Probabilistic Tractography with crossing fibers (ProbTrackX) available with FSL 6.0 (Hernandez-Fernandez et al., 2019), and the default (one-way) setting with a loop check. The resulting output describe the density of WM connections between the ROI pair.

To improve the tractography results, and to reduce the incidence of erroneous streamlines, we employed exclusion masks. These exclusion masks were based on a mask of CSF created by thresholding the MD maps (voxels with MD >2.10^−3^mm^2^/s were considered as CSF) and corrected by removing voxels with FA >0.25 (which might correspond to WM voxels in the corpus callosum but close to the ventricles). The exclusion masks were further adapted to exclude all other brain structures apart from the considered ROIs pair. Additionally, where the pair of ROIs were ipsilateral, i.e., in a single hemisphere, the entire contralateral hemisphere was also excluded. No supplementary constraints were included in the tractography runs.

Reconstructed tracts were then thresholded at 5% of the maximum fiber density of the evaluated tract. This was not performed for cortico-cortical inter-hemispheric tracts, whose reconstructions were used in their original state due to low streamline numbers.

The final list of SM tracts of interest (corresponding to homotopic inter-hemispheric tracts, short-range S1-M1 intra-hemispheric tracts, and long-range intra-hemispheric cortico-subcortical tracts) is described in **Figure**

**1.B**. Note that the (inter- and intra-hemispheric) cortico-cortical tracts involving paracentral regions could not be evaluated due to frequent tractography errors identified upon visual examination.

#### Extraction of tract-specific metrics

DTI and NODDI-derived metrics (FA, MD, AD, RD, NDI, ODI) were extracted from each individual tract by calculating the weighted average value (metric 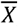) using the following equation:

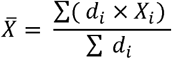

where *i* denotes the tract voxels, *d*_*i*_ is the fiber density at voxel *i* of a tract, and *X*_*i*_ is value of the metric at voxel *i* (Hua et al., 2008). This weighted approach gives more weight to the central portion (with higher fibre density) compared to the tract periphery, renders the measures independent on the number of streamlines assessed by the tractography algorithm, and limits the effect of potential artefacts related to tractography reconstruction.

### Statistical analyses

#### Descriptive models and univariate analyses

Considering all tracts except interhemispheric connections, we first assessed whether different factors (namely the tract, the hemisphere, and the infant group) have an effect on each diffusion metric by implementing a global ANOVA model in R (version 4.0.5, 2021.03.31). Although the hemisphere factor appeared to have a significant effect for some metrics (**Table SupT3.A**), we decided to average diffusion metrics for the left and right tracts in all subsequent analyses to explore group differences, as no significant interaction was observed between hemisphere and group for any of the 6 evaluated metrics. The differences between groups of interest (PT_EV_ vs FT_EVCt_; PT_ML_ vs FT_MLCt_) was further studied with paired t-tests (over all tracts) corrected for multiple comparisons across the metrics.

In a second step, we aimed to select clinical factors (focusing on the ones previously reported as having significant effect on infants WM microstructure, cf. section ***Neonatal characteristics at birth***) with a relevant effect on the diffusion metrics in our cohort. We thus performed additional ANOVA modelling considering all tracts (including interhemispheric) on the PT group, as detailed clinical variables were present only in this group (**Table SupT5**). This model allowed identification of confounders that should be taken into account in the following analyses of the diffusion metrics.

We then performed a complete global ANCOVA model on each metric for all tracts together with the significant clinical factors identified in the previous steps as well as additional *continuous* variables: GA at birth (instead of group categorization), PMA at scan and a proxy of whole-brain WM microstructure to account for additional interindividual variability. For this latter variable (hereafter named *WM residuals)*, we considered the *residuals* of the linear model considering the metric averaged over the whole WM mask as a function of *GA at birth and PMA at scan*, which were shown to be highly associated with the averaged WM metric (**Table SupT6.A**). This final model allowed us to identify the confounders to be taken into account in all the subsequent analyses.

#### Univariate tract-specific analyses

Finally, to investigate tract-specific univariate relationships between the diffusion metrics and GA at birth, we performed a linear regression of each metric per tract, adjusted for PMA at scan and WM residuals (the only confounders shown to be relevant in the previous analyses). The slope tests of the linear fits were corrected for multiple comparisons across all studied metrics and tracts. Additionally, we estimated a proxy of the maturational gap of the PT_EV_ and PT_ML_ groups by calculating relative percentage changes in the metric values between the PT_EV_ and PT_ML_ compared to their respective matched FT controls.

### Multivariate tract-specific analyses on effects of prematurity

In order to characterize the potential difference present at TEA between the microstructural profiles of PT infants compared to FT infants for each tract, we used a previously proposed multiparametric approach (Kulikova et al., 2015) based on the Mahalanobis distance. The goal was to evaluate the distance between each individual PT infant and the FT group as a reference, by taking into account the inter-subject variability within the FT group and the collinearity between a set of diffusion metrics.

Firstly, we scaled each diffusion metric between [0; 1], considering all tracts and the mean WM in all the PT and FT infants. The tract scaled metrics were then corrected for GA at birth, PMA at scan and WM residuals (cf. section ***Descriptive models and univariate analyses***), considering each of the 3 groups independently (PT_EV_, PT_ML_, and entire FT group) and keeping the respective group value means. Next, we divided the PT and FT *individual* tract metric values by their respective metric means from the FT group.

For the calculation of the Mahalanobis distance, it is assumed to be beneficial to choose independent metrics that provide complementary information. With this in mind, we decided to subset the six metrics into 3 parallel analysis streams based on the nature of the metrics and models used to derive them. AD and RD, which are direct measures of the diffusivity within the tracts were retained as *set 1*. More complex but commonly used DTI metrics: MD and FA, formed *set 2*. Finally, NODDI metrics (NDI and ODI) formed an independent *set 3* to dissociate them from the more widely established DTI metrics and test their relevance for microstructure in the context of SM network and prematurity.

For a given tract, the Mahalanobis distance (Dtract) for a given PT individual was then computed using the following equation:

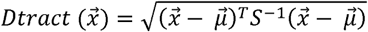

Where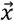 is a multivariate vector describing the PT individual tract-specific metrics,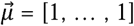 is the mean vector for the corresponding FT group, and S is a covariation matrix for diffusion metrics in FT infants.

In the interpretation, the smaller the distance, the closer the individual preterm infant is to the distribution within the control FT cohort. Differences in distances across tracts can be interpreted as a differential, tract-specific effect of prematurity on maturation.

First, we evaluated whether the distances for each of the two PT subgroups were significantly different from 0 (meaning that the PT subgroups are different from the FT reference group) using one-sample Wilcoxon signed rank tests corrected for multiple comparisons across all tracts and metric sets. Similarly, to the previous descriptive analysis, we further confirmed the effect of tracts and PT subgroups on the Mahalanobis distances using global ANOVA modelling. We additionally compared distances between the two PT subgroups based on unpaired t-tests corrected for multiple comparisons (considering all tracts or each tract separately). Both ANOVA and t-tests were performed after checking for normality of the Mahalanobis bundle values using Shapiro-Wilk test corrected for multiple comparisons across sets. To establish whether SM tracts were differentially affected by the prematurity status, we compared all possible pairs of tracts within each PT group, using paired t-tests corrected for multiple comparisons across all studied metric sets and tracts.

We finally aimed to evaluate whether this maturational distance related to prematurity in each PT subgroup was related to later neurodevelopmental outcome. For each tract, we studied Pearson’ s correlations between the Mahalanobis metrics sets and the 5 BSID-III scaled scores, considering infants with outcome data in each group separately (PT_EV_ N=24; PT_ML_ N=20). The reported results were corrected for multiple comparisons across all tract and metric sets. Statistical tests throughout the analyses were considered with a 0.95 significance level.

## Results

### Cohort characteristics

A summary of characteristics for each group and detailed results of the group comparisons are presented in **Table 1**. Obstetric factors, multiple pregnancies, IUGR, and delivery by caesarean section were significantly more frequent in the PT group compared to FT (30.5% vs 1.5%, 31% vs 2%, and 68% vs 49% respectively). As expected, PT infants differed significantly from FT group in weight, length, and head circumference at birth, as well as Apgar scores. Among the neonates admitted to NICU, only PT needed surfactant, ventilatory support and parenteral nutrition, with 7 (13%) infants needing mechanical ventilation >7 days and 4 (7.5%) parenteral nutrition >21 days.

**Table 1.**
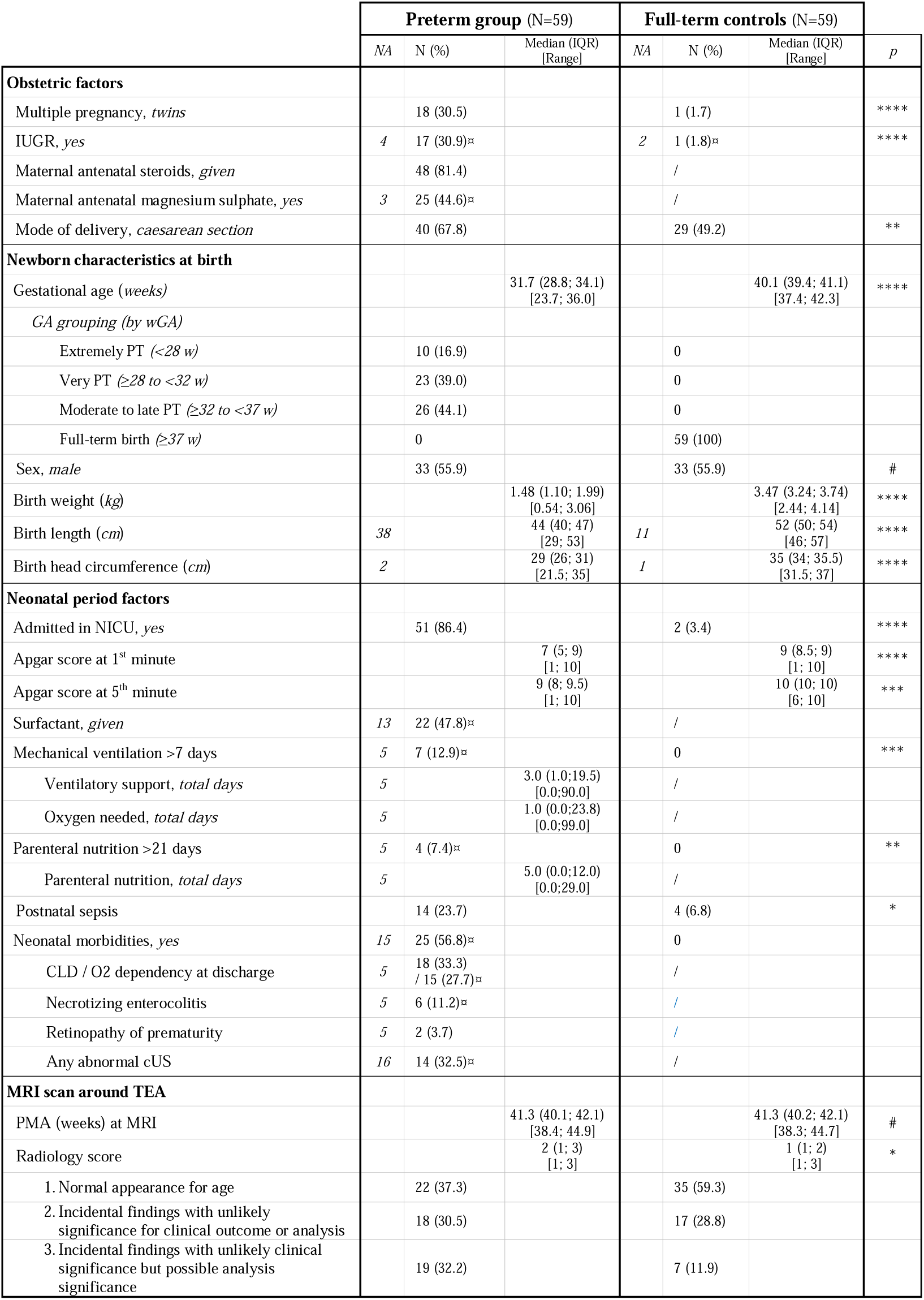
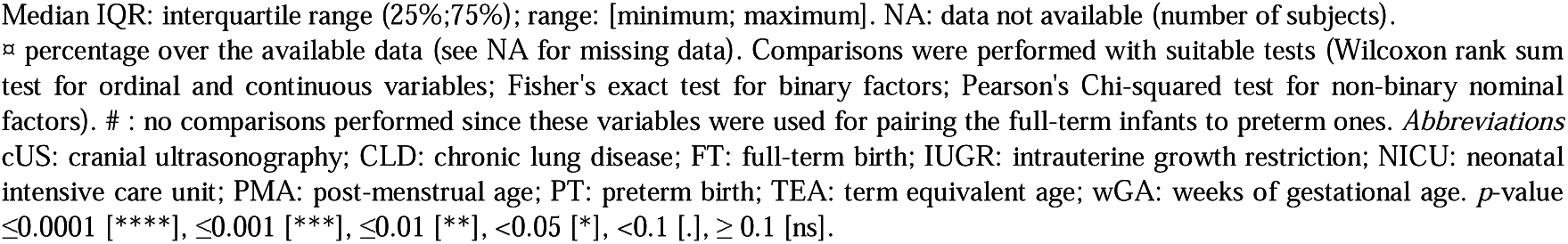
Detailed clinical and sociodemographic information for the 59 pairs of infants.

|  | Preterm group (N=59) |  |  | Full-term controls (N=59) |  |  | *p* |
| --- | --- | --- | --- | --- | --- | --- | --- |
|  | *NA* | N (%) | Median (IQR)<br>[Range] | *NA* | N (%) | Median (IQR)<br>[Range] |  |
| **Obstetric factors** |  |  |  |  |  |  |  |
| Multiple pregnancy, *twins* |  | 18 (30.5) |  |  | 1 (1.7) |  | **** |
| IUGR, *yes* | 4 | 17 (30.9)▯ |  | 2 | 1 (1.8)▯ |  | **** |
| Maternal antenatal steroids, *given* |  | 48 (81.4) |  |  | / |  |  |
| Maternal antenatal magnesium sulphate, *yes* | 3 | 25 (44.6)▯ |  |  | / |  |  |
| Mode of delivery, *caesarean section* |  | 40 (67.8) |  |  | 29 (49.2) |  | ** |
| **Newborn characteristics at birth** |  |  |  |  |  |  |  |
| Gestational age (*weeks*) |  |  | 31.7 (28.8; 34.1)<br>[23.7; 36.0] |  |  | 40.1 (39.4; 41.1)<br>[37.4; 42.3] | **** |
| *GA grouping (by wGA)* |  |  |  |  |  |  |  |
| Extremely PT (<28 *w*) |  | 10 (16.9) |  |  | 0 |  |  |
| Very PT (≥28 *to* <32 *w*) |  | 23 (39.0) |  |  | 0 |  |  |
| Moderate to late PT (≥32 *to* <37 *w*) |  | 26 (44.1) |  |  | 0 |  |  |
| Full-term birth (≥37 *w*) |  | 0 |  |  | 59 (100) |  |  |
| Sex, *male* |  | 33 (55.9) |  |  | 33 (55.9) |  | # |
| Birth weight (*kg*) |  |  | 1.48 (1.10; 1.99)<br>[0.54; 3.06] |  |  | 3.47 (3.24; 3.74)<br>[2.44; 4.14] | **** |
| Birth length (*cm*) | 38 |  | 44 (40; 47)<br>[29; 53] | 11 |  | 52 (50; 54)<br>[46; 57] | **** |
| Birth head circumference (*cm*) | 2 |  | 29 (26; 31)<br>[21.5; 35] | 1 |  | 35 (34; 35.5)<br>[31.5; 37] | **** |
| **Neonatal period factors** |  |  |  |  |  |  |  |
| Admitted in NICU, *yes* |  | 51 (86.4) |  |  | 2 (3.4) |  | **** |
| Apgar score at 1<sup>st</sup> minute |  |  | 7 (5; 9)<br>[1; 10] |  |  | 9 (8.5; 9)<br>[1; 10] | **** |
| Apgar score at 5<sup>th</sup> minute |  |  | 9 (8; 9.5)<br>[1; 10] |  |  | 10 (10; 10)<br>[6; 10] | *** |
| Surfactant, *given* | 13 | 22 (47.8)▯ |  |  | / |  |  |
| Mechanical ventilation >7 days | 5 | 7 (12.9)▯ |  |  | 0 |  | *** |
| Ventilatory support, *total days* | 5 |  | 3.0 (1.0;19.5)<br>[0.0;90.0] |  | / |  |  |
| Oxygen needed, *total days* | 5 |  | 1.0 (0.0;23.8)<br>[0.0;99.0] |  | / |  |  |
| Parenteral nutrition >21 days | 5 | 4 (7.4)▯ |  |  | 0 |  | ** |
| Parenteral nutrition, *total days* | 5 |  | 5.0 (0.0;12.0)<br>[0.0;29.0] |  | / |  |  |
| Postnatal sepsis |  | 14 (23.7) |  |  | 4 (6.8) |  | * |
| Neonatal morbidities, *yes* | 15 | 25 (56.8)▯ |  |  | 0 |  |  |
| CLD / O2 dependency at discharge | 5 | 18 (33.3)<br>/ 15 (27.7)▯ |  |  | / |  |  |
| Necrotizing enterocolitis | 5 | 6 (11.2)▯ |  |  | / |  |  |
| Retinopathy of prematurity | 5 | 2 (3.7) |  |  | / |  |  |
| Any abnormal cUS | 16 | 14 (32.5)▯ |  |  | / |  |  |
| **MRI scan around TEA** |  |  |  |  |  |  |  |
| PMA (weeks) at MRI |  |  | 41.3 (40.1; 42.1)<br>[38.4; 44.9] |  |  | 41.3 (40.2; 42.1)<br>[38.3; 44.7] | # |
| Radiology score |  |  | 2 (1; 3)<br>[1; 3] |  |  | 1 (1; 2)<br>[1; 3] | * |
| 1. Normal appearance for age |  | 22 (37.3) |  |  | 35 (59.3) |  |  |
| 2. Incidental findings with unlikely significance for clinical outcome or analysis |  | 18 (30.5) |  |  | 17 (28.8) |  |  |
| 3. Incidental findings with unlikely clinical significance but possible analysis significance |  | 19 (32.2) |  |  | 7 (11.9) |  |  |

Median IQR: interquartile range (25%;75%); range: [minimum; maximum]. NA: data not available (number of subjects).
⌘ percentage over the available data (see NA for missing data). Comparisons were performed with suitable tests (Wilcoxon rank sum test for ordinal and continuous variables; Fisher's exact test for binary factors; Pearson's Chi-squared test for non-binary nominal factors). # : no comparisons performed since these variables were used for pairing the full-term infants to preterm ones. *Abbreviations* cUS: cranial ultrasonography; CLD: chronic lung disease; FT: full-term birth; IUGR: intrauterine growth restriction; NICU: neonatal intensive care unit; PMA: post-menstrual age; PT: preterm birth; TEA: term equivalent age; wGA: weeks of gestational age. *p*-value $\leq 0.0001$ [\*\*\*\*], $\leq 0.001$ [\*\*\*], $\leq 0.01$ [\*\*], $< 0.05$ [\*], $< 0.1$ [·], $\geq 0.1$ [ns].

Morbidities linked to prematurity were seen in 25 PT infants (56.8% of the 44 with available data), including chronic lung disease for 18 infants (15 needing oxygen at discharge), 16 infants had an abnormality identified on cUS during NICU period, 6 had NEC and 2 had ROP.

As would be expected, differences were also observed in Radiology scores between PT and FT babies, with more PT infants having a score equal to 3. Nevertheless, as scores of 3 or less are considered to be of unlikely clinical significance, we did not further analyze this variable.

Selected clinical descriptors for the 4 infant subgroups included in the next descriptive analyses are presented in **Table SupT1**.

### Neurodevelopmental outcome and characteristics at 18 months

Significant differences were observed in IMD scores reported at 18mCA (n=96) (**Table SupT2**), with PT families tending to live in more deprived areas than the FT ones (quintiles ≤3 for 64.3% vs 29.6% respectively, *data not shown*).

Among the 97 infants with available BSID-III data, no significant differences between PT and FT controls were observed for the corrected age at assessment. Paired t-tests on scaled scores showed no significant differences between PT and FT groups after correcting the results for multiple comparisons (**Figure 2**). BSID- III scaled scores across PT_EV_, PT_ML_ FT_EVCt_ and FT_MLCt_ subgroups (presented in **Figure SupF1**) also showed no significant group effect.

**Figure 2.**
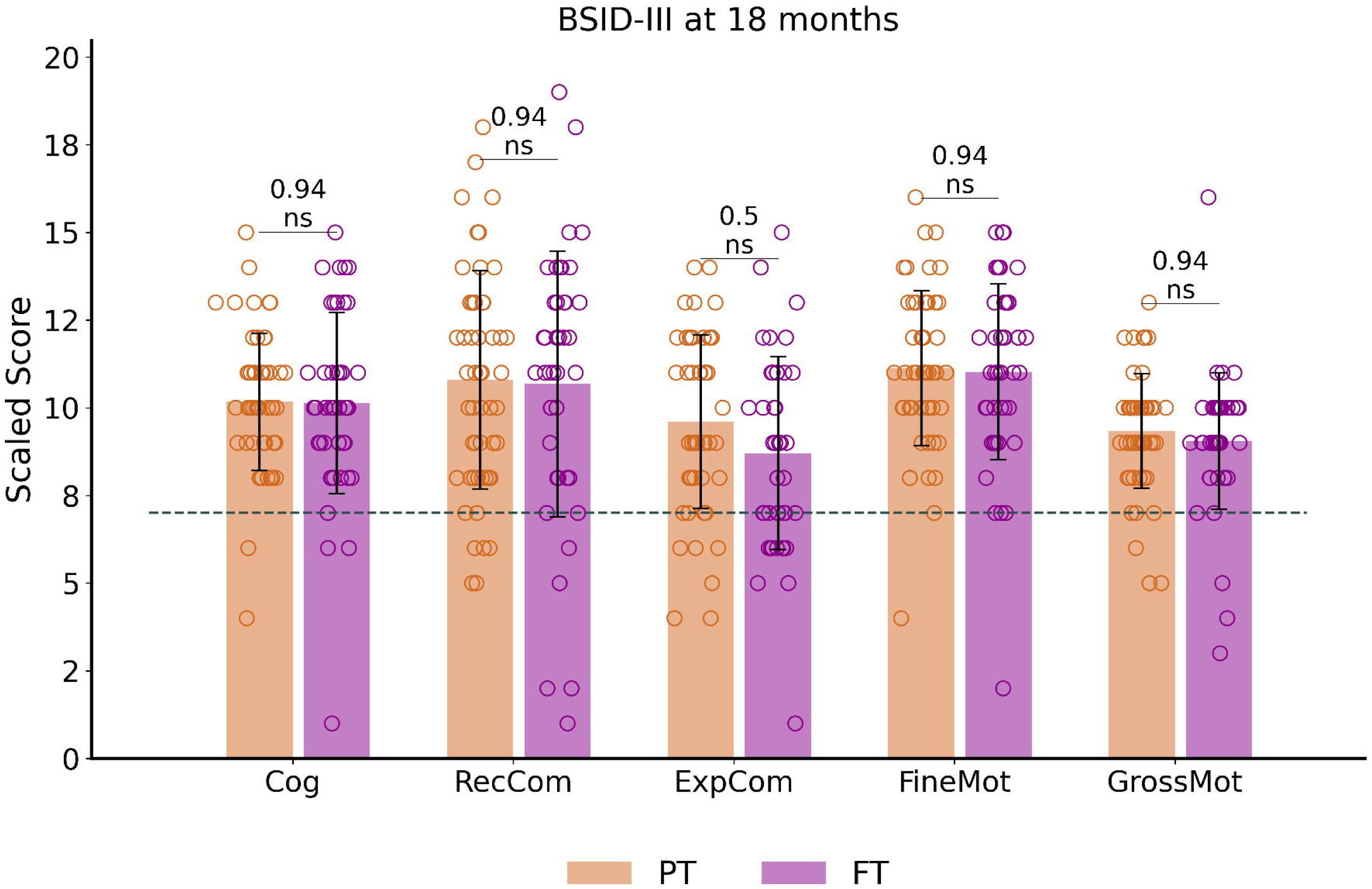
Outcome assessment at around 18 months of corrected age: BSID-III scaled scores distribution and comparisons between preterm and full-term infant groups. The dotted line corresponds to the pathological threshold, with scores < 7 (< -1 SD) indicating a developmental delay. Of note, only one extreme PT (male, born at 27.6wGA) presented severe developmental delay (scores <-3SD) for Cognitive and both Communication scores (with Fine motor score at –2SD and Gross motor score on the norm values). Reported *p*-values come from t-tests corrected for multiple comparisons. See *Supplementary Figure SupF1* for infant subgroup analysis using one-way ANOVA. BSID-III: Bayley Scales of Infant and toddler Development, 3rd edition. PT: preterm, FT: full-term. Cog: cognitive; RecCom: receptive communication, ExpCom: expressive communication; FineMot: fine motor, GrossMot: gross motor scaled scores. ns: non significant.

Over the whole cohort, only a small number of infants obtained scaled scores indicating a developmental delay (scores below 7 and corresponding to <-1SD), with no significant difference between PT and FT. These consisted of developmental delay in 30.9% of infants for expressive communication (N=30, 19 PT), 14.4% for receptive communication (N=14, 7 PT), 11.3% for gross motor (N=11, 5 PT), 6.2% for fine motor (N=6, 4 PT) and cognition (N=6, 4 PT).

### Description of SM tract reconstructions

Visual inspection of the automated reconstructions for all tracts was performed on 12 randomly selected infants which allowed us to evaluate the quality of reconstructions for all the 15 tracts of interest, in a similar way across PT and FT infants. Examples of individual tract reconstructions and diffusion metric maps are shown in **Figure 1 (C-D)** for a representative FT infant.

### Descriptive models

#### Differences in diffusion metrics across tracts and groups

The initial ANOVA examined the influence of the tract, hemisphere and infant group on each diffusion metric (**Table SupT3.A**). It confirmed the expected significant effects of tract and group, as well as the interaction between both, except for FA where the group effect did not reach the significance level. A significant effect of hemisphere was further observed for FA, RD and ODI metrics as well as interaction between tract and hemisphere, uncovering asymmetries between left and right tracts. However, given that no significant interaction was observed between hemisphere and group for any of the six metrics, we decided not to further explore these differences between left and right tracts, and to consider *averaged diffusion metrics for left and right tracts* in all subsequent analyses. Tract-specific distributions of diffusion metrics across the 4 groups are presented in **Figure 3**. Post-hoc group analyses (paired t-tests considering all tracts) corroborated the existence of significant differences between PT infants and their matched FT controls, with higher AD, RD and MD metrics, opposed to lower FA, NDI, and ODI metrics in PT groups compared to FT, especially in the PT_EV_ vs FT_EVCt_ (**Table SupT3.B** and **Figure 3**). This suggested that the more preterm the infant is, the more “immature” microstructural characteristics are. Group comparisons for PT_EV_ vs FT_EVCt_ are presented per tract in **Table SupT4** (group comparisons between PT_ML_ vs FT_MLCt_ were not significant).

**Figure 3.**
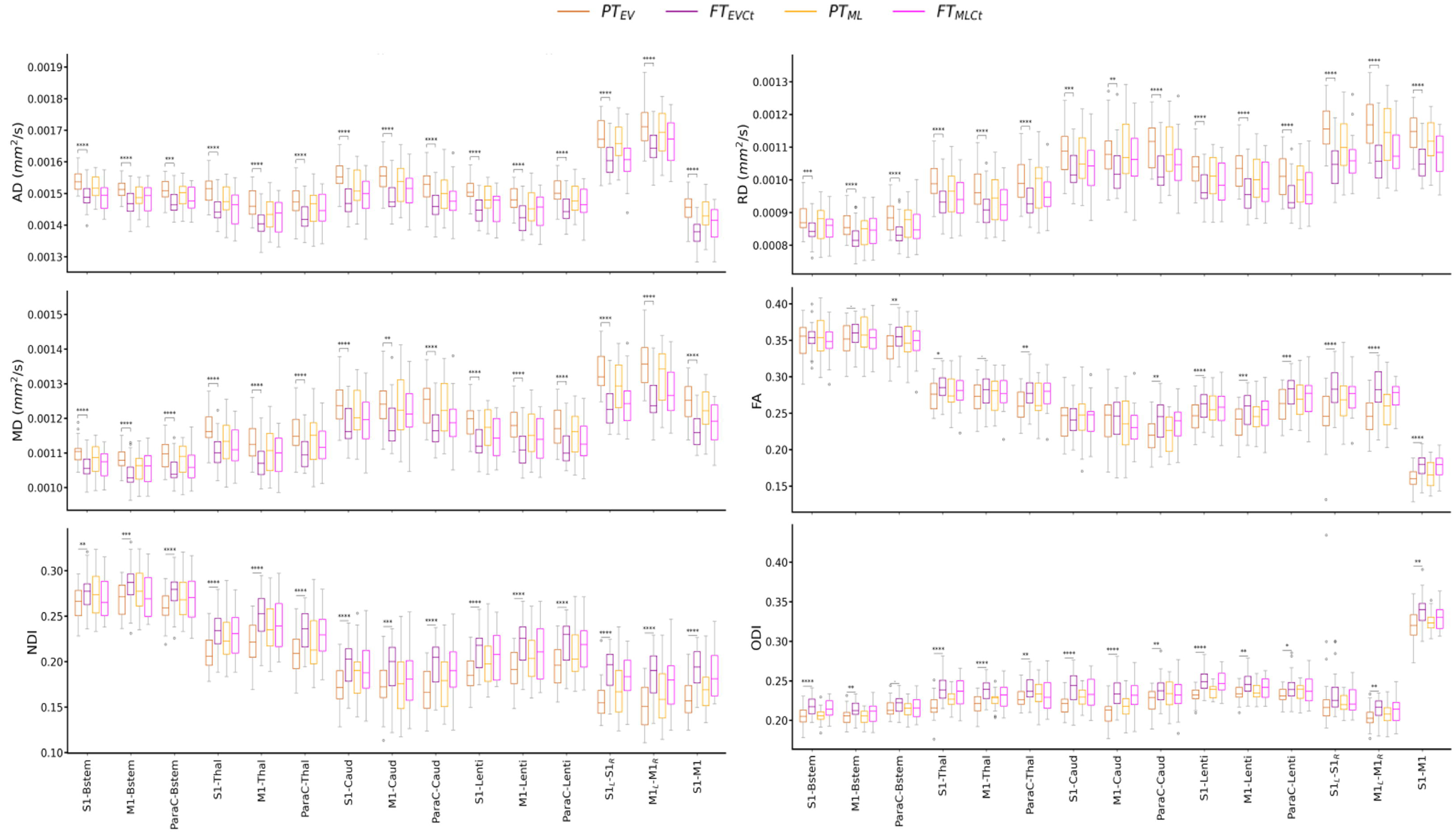
Diffusion metrics across tracts and cohort subgroups: *extreme to very preterm* group (PT_EV_, dark orange) compared to *paired full-term controls* (FT_EVCt_, dark purple), and *moderate to late preterm* group (PT_ML_, light orange) compared to *paired controls* (FT_MLCt_, light purple). Significances are results of the tract-specific paired t-tests between paired groups, corrected for multiple comparisons. Only the comparisons between PT_EV_ – FT_EVCt_ reached significance. Refer to *Figure 1* legend for abbreviations and to *Table 1* for *p*-value legend.

Visual assessment of the metric distributions (**Figure 3**) revealed important microstructural differences between tracts, and suggested some similarities across metrics across all groups (e.g., RD-MD having opposite pattern to NDI across all tracts).

#### Effect of clinical variables on diffusion metrics

The second ANOVA, focusing on PT infants, allowed study of the effect of selected clinical factors on the diffusion metrics and to identify potential confounders for the subsequent comparisons (**Table SupT5**). Confirming previous results, the tract effect was highly significant for all metrics. The group effect as well as the interaction between the tract and group, were less significant than in the previous analyses, probably because only PT infants were considered while larger differences are observed between PT and FT than between PT_EV_ and PT_ML_. Globally, multiple pregnancy, IUGR and parenteral nutrition were weakly associated with some of the evaluated diffusion metrics. Interestingly, sex and preterm morbidities were not significantly associated with the diffusion metrics, which led us not to retain this latter variable in the next analyses.

#### Global modelling of the diffusion metrics

In a complementary analysis evaluating the whole cohort, the effects of supplementary *continuous* variables: GA at birth (instead of group belonging), PMA at scan, and WM residuals (**Table SupT6.A**) were added to the clinical factors selected in the previous analysis. For each metric, the modelling uncovered significant associations with the three variables, but the effects of sex and clinical factors, other than IUGR, dropped considerably in significance compared to the previous models probably because these factors are related to GA at birth and WM residuals, leading us not to retain them for the next analyses. Moreover, as the effect of IUGR was negligible compared to the effects of GA at birth, PMA at scan and WM residuals, this factor was likewise not retained in the subsequent analyses.

### Changes of tract-specific diffusion metrics with GA at birth

To further explore changes in diffusion metrics with GA at birth, we performed a linear regression over the whole cohort after correction for significant variables identified in the previous analyses (PMA at scan and WM residuals). Significant associations with GA at birth were observed in almost all SM tracts and metrics (**Table 2, Table SupT6.B** and **Figure SupF2**). Confirming the observations from the group analyses (**Figure 3, Table SupT4**), MD, AD, and RD metrics decreased, and FA, NDI, and ODI increased with the GA at birth. The lower associations for FA (only 10/15 tracts) and ODI (13/15 tracts) (**Table 2**) suggested that these metrics might be less sensitive to detect the variation of microstructure characteristics with GA at birth within some SM tracts, at least at TEA.

**Table 2.** Univariate linear regression analysis: relationship between SM tracts’ diffusion metrics (corrected for PMA at scan and WM residuals) and GA at birth over the whole cohort.

| Tracts | AD |  | RD |  | MD |  | FA |  | NDI |  | ODI |  |
| --- | --- | --- | --- | --- | --- | --- | --- | --- | --- | --- | --- | --- |
|  | T | *p* | T | *p* | T | *p* | T | *p* | T | *p* | T | *p* |
| S1-Bstem | -7.86 | **** | -4.83 | **** | -7.32 | **** | -0.33 | ns | 2.98 | ** | 6.28 | **** |
| M1-Bstem | -6.43 | **** | -5.22 | **** | -6.84 | **** | 0.69 | ns | 3.41 | ** | 3.74 | *** |
| ParaC-Bstem | -6.74 | **** | -7.58 | **** | -8.88 | **** | 2.89 | ** | 5.57 | **** | 1.79 | ns |
| S1-Thal | -9.88 | **** | -8.72 | **** | -9.81 | **** | 3.14 | ** | 8.29 | **** | 7.74 | **** |
| M1-Thal | -7.7 | **** | -7.22 | **** | -8.19 | **** | 1.89 | ns | 7.14 | **** | 5.24 | **** |
| ParaC-Thal | -5.76 | **** | -8.05 | **** | -7.87 | **** | 4.89 | *** | 7.53 | **** | 2.26 | * |
| S1-Caud | -11.1 | **** | -6.1 | **** | -8.19 | **** | 0.36 | ns | 6.82 | **** | 6.45 | **** |
| M1-Caud | -7.82 | **** | -3.04 | ** | -4.41 | **** | -1.19 | ns | 4.4 | **** | 7.3 | **** |
| ParaC-Caud | -6.42 | **** | -9.52 | **** | -9.21 | **** | 5.49 | *** | 9.34 | **** | 3.28 | ** |
| S1-Lenti | -10.2 | **** | -10.8 | **** | -11.7 | **** | 5.1 | *** | 10.2 | **** | 9.3 | **** |
| M1-Lenti | -7.87 | **** | -9.96 | **** | -10.1 | **** | 5.09 | *** | 9.49 | **** | 4.35 | **** |
| ParaC-Lenti | -8.2 | **** | -10.21 | **** | -10.9 | **** | 4.49 | *** | 9.26 | **** | 3.09 | ** |
| S1L-S1R | -7.22 | **** | -12.1 | **** | -12.6 | **** | 7.07 | *** | 9.16 | **** | 1.35 | ns |
| M1L-M1R | -6.62 | **** | -12.58 | **** | -11.8 | **** | 9.07 | *** | 10.58 | **** | 4.08 | *** |
| S1-M1 | -11.5 | **** | -15.68 | **** | -14.7 | **** | 9.41 | *** | 13.04 | **** | 4.93 | **** |

We further analyzed the relative percent difference in diffusion metrics between the PT_EV_ and PT_ML_ infants compared to their paired FT neonates, in order to estimate a proxy of the maturational gap related to prematurity (**Figure SupF3**). Visual inspection suggested a larger gap in the PT_EV_ subgroup than in PT_ML_, highlighting the effect of prematurity degree on some tract’ s microstructural characteristics. However, the observed variability between the metrics rendered the interpretation of different maturational patterns across tracts difficult, justifying the need for a multivariate approach.

### Mahalanobis distance of PT subjects from the typical FT profile

To explore the impact of prematurity on the tract-specific microstructure, we computed multi-metric Mahalanobis distances of PT subgroups (PT_EV_ and PT_ML_ independently) to all FT infants as reference, using the 3 metric sets: *set 1* (AD and RD), *set 2* (MD and FA), and *set 3* (NDI and ODI) (**Figure 4**). For a given tract, computed Mahalanobis distance can be understood as a ***maturational distance*** for a given PT infant compared to the FT group.

**Figure 4.**
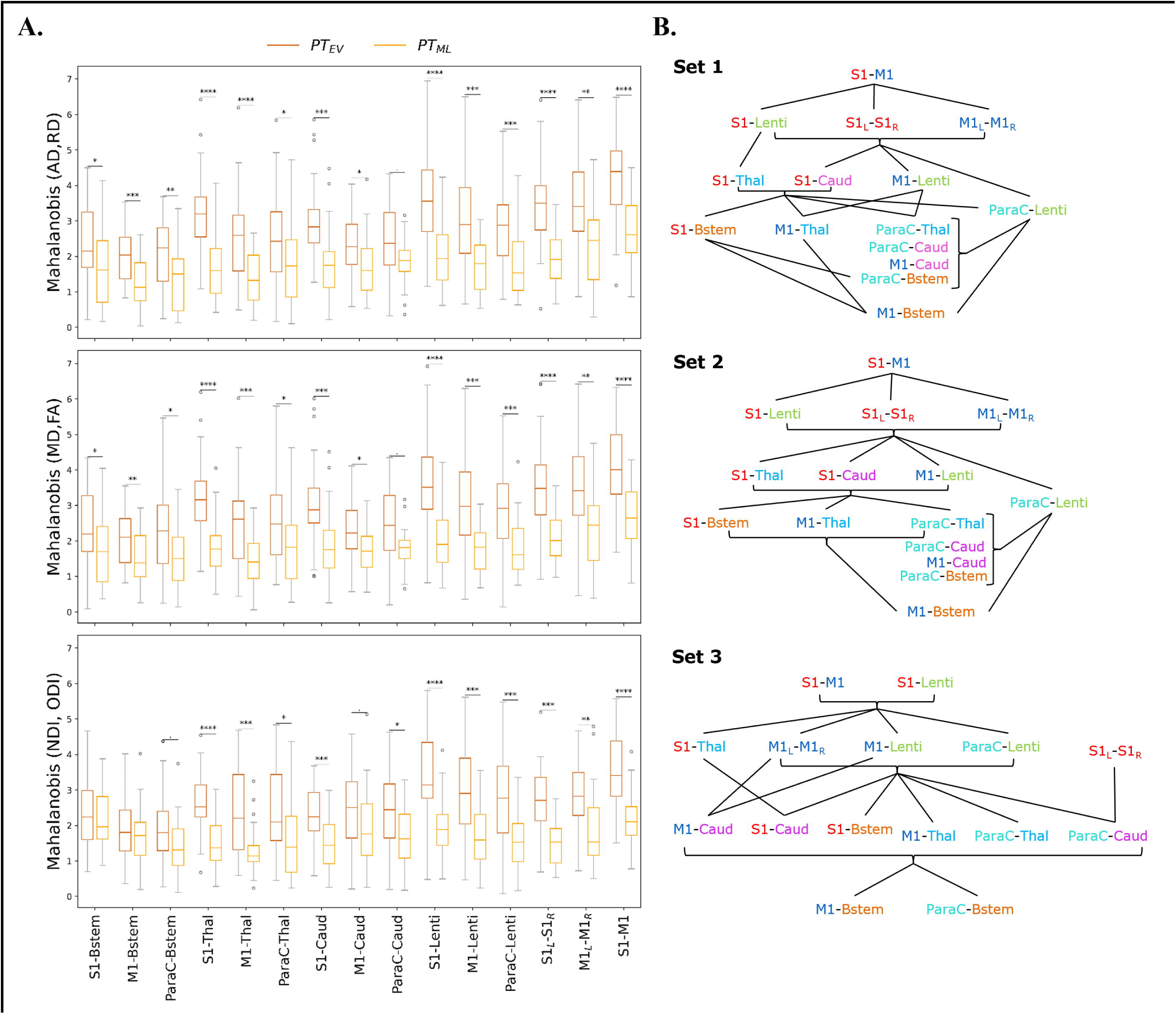
A. Multi-metric Mahalanobis distance across tracts and PT subgroups. (PT_EV_ and PT_ML_ with FT controls as reference) at TEA. The smaller the distance, the less the microstructural profile of the PT infant differs from the FT reference group. Note that the effect of prematurity is globally smaller for PT_ML_ than for PT_EV_ infants across the studied SM tracts. Significances are results of the comparison of Mahalanobis distances (PT_EV_ vs PT_ML_) with unpaired t-tests (p-values corrected for multiple corrections) for each set and each tract. For visualization purposes, outliers (mean ±3SD) were removed (6 points for PT_EV_, 3 for PT_ML_ infants). **B. Order of the SM tracts in the PT**_**EV**_ **subgroup based on the Mahalanobis distance per metrics set** (higher values on the top), highlighting the differential effect of prematurity on SM tracts microstructure. The lines represent the significant differences between tracts according to paired t-tests (corrected for multiple comparisons) over the PT_EV_ group (for visualisation purposes, the statistical threshold was relaxed to p<0.1). Metrics sets: 1 (AD, RD); 2 (MD, FA); 3 (NDI, ODI). Refer to *Figures 1* legend for ROIs color code and abbreviations.

For both PT subgroups (PT_EV_ and PT_ML_), all sets and all tracts, distances were highly significantly different from 0 as assessed by Wilcoxon tests corrected for multiple comparisons (all p<0.001), suggesting that SM network microstructure as estimated with this approach is affected by prematurity, even moderate or late. Considering all PT infants, ANOVA modelling on distances for each set confirmed the expected effects of group, tract, and the interaction between group and tract for the three metric sets (**Table SupT7**). As expected, the distance increased with the prematurity levels, with unpaired t-tests per tract comparing the two PT subgroups revealing higher distances in PT_EV_ than in PT_ML_ (**Figure 4.A**). In addition, the tracts were not affected in the same manner: distances were different between PT_EV_ and PT_ML_ for all tracts except for ParaC- Caud in both DTI sets (1 and 2), and for set 3: S1-Bstem, M1-Bstem, ParaC-Bstem and M1-Caud.

#### Tract-specific effects of prematurity

To further evaluate the differential effect of prematurity on specific tracts, we subsequently compared each pair of tracts through paired t-tests in each PT subgroup independently (**Figure SupF4**). While many more significant tract-by-tract differences were observed in the PT_EV_ than in the PT_ML_ group (69 vs 26/105 for *set 1*; 70 vs 22/105 for *set 2*; 61 vs 23/105 for *set 3*), the results were rather consistent between the two PT subgroups, with only a few tract-by-tract differences observed in the PT_ML_ group but not in the PT_EV_ group (3/105 for *set 1*; 2/105 for *set 2*; 7/105 for *set 3*).

Focusing on the **PT**_**EV**_ **subgroup**, the significant differences between tracts assessed by the paired t-tests allowed to propose an ordering of the tracts based on the relative effects of prematurity on microstructural characteristics (**Figure 4.B**). For *sets 1 and 2* (DTI sets), the orderings were highly similar, with, somewhat schematically, the following tracts showing the lowest to highest distances: 1) M1-Bstem; 2) ParaC-Bstem, S1-Bstem, M1-Thal, ParaC-Thal, M1-Caud, ParaC-Caud; 3) M1-Lenti, ParaC-Lenti, S1-Thal, S1-Caud; 4) S1- Lenti, S1L-S1R, M1L-M1R; 5) S1-M1. For *set 3* (NODDI set), the ordering showed schematically, from the lowest to highest distances of tracts: 1) M1-Bstem, ParaC-Bstem; 2) S1-Bstem, M1-Thal, ParaC-Thal, M1- Caud, ParaC-Caud, S1-Caud; 3) S1-Thal, M1-Lenti, ParaC-Lenti, S1L-S1R, M1L-M1R; 4) S1-Lenti, S1-M1 (**Figure 4.B**).

Despite a few differences in the ordering of a couple of tracts, results were quite consistent across the three sets and revealed a differential impact of prematurity on the SM tracts microstructure. Overall, the tract ordering based on maturational distances highlighted a coherent **caudo-rostral and central-to-periphery pattern**, with: the cortico**-Brainstem** tracts presenting the lowest distances and thus the least impact of prematurity; the cortico-**Thalamic** and cortico-**Caudate** tracts showing intermediate distances; the cortico- **Lenticular** tracts appearing with the highest distances among the cortico-subcortical tracts; and the **cortico-cortical** tracts revealing the highest impact of prematurity, but again with a gradation since inter-hemispheric tracts (S1L-S1R and M1L-M1R) showed lower distances than the intra-hemispheric tracts (S1-M1). Interestingly, the **S1 tracts** presented, in an almost systematic way, higher distances than the M1 and ParaCentral tracts, both presenting similar profiles.

This approach of tract ordering based on the maturational distance related to prematurity was not considered for the **PT**_**ML**_ **subgroup** as tract pairwise comparisons were less systematically significant than in the PT_EV_ group and the ordering was more difficult to synthesize. For this subgroup, the distances of all tracts were more homogeneous (**Figure SupF4**), which may be associated with a lesser effect of prematurity on the microstructural profiles of the SM tracts.

#### Tract-specific maturational distance associated with neurodevelopmental outcome

Finally, we assessed whether maturational distances related to prematurity at TEA might be related to outcome (BSID-III scaled scores) at 18mCA, considering each PT subgroup independently. Pearson correlations showed significant results only in **PT**_**EV**_ group (N=24), for *Set 3* (**NODDI**) (**Table SupT8**) and for specific tracts: *M1-Bstem and ParaC-Bstem* distances were both negatively correlated with *Cognitive* scaled score and *Fine motor* score, while *M1-Lenti, ParaC-Lenti* and *S1-M1* tracts were also negatively correlated with *Fine motor* score (the lower the maturational distance, the higher the score and thus the better the outcome). Interestingly, these 5 tracts showed different levels of distances over the PT_EV_ group, with M1- and ParaC- Bstem having a distance closest to 0, whereas S1-M1 was the tract with the highest distance; and M1- and ParaC-Lenti presented a similar and intermediate distance.

## Discussion

In this study, we observed significant differences in the microstructure of most SM tracts between low-risk PT and FT infants at TEA, based on univariate analyses of DTI and NODDI metrics. Moreover, a multi- parametric assessment suggested that the maturational distance between these infants differ with GA at birth and across SM tracts, with alterations affecting in particular the S1-related tracts and the more rostral tracts (i.e., the cortico-lenticular ones). Correlations between the NODDI maturational distance of specific tracts and BSID-III scaled scores evaluated at 18mCA were also observed.

### I. Exploring the developing SM network with diffusion MRI

#### 1. Identifying SM structures of interest in infants

Somatosensory and motor systems in the mature brains consist of distributed and interconnected cortical and subcortical regions. In our study, we disregarded *non-primary cortical areas* (supposed to mature later during typical development) focusing instead on the primary core of cortical and deep GM structures (S1, M1, thalamus, BG, brainstem). These structures interact through cortico-cortical connections but also indirectly through cortico-subcortical loops that develop early and are supposed to underpin a wide range of early SM experiences (including the diverse fetal and neonate perceptions and spontaneous motor behaviors).

When focusing on primary SM structures in the early developmental context, **S1** and **M1** areas appear essential for processing peripheral somatosensory inputs, initiating and controlling motor behaviors. Proper functioning of the BG appears necessary for the motor cortex to relay appropriate motor commands to the lower levels of the hierarchy and finally to the spinal cord and peripheral nervous system through connections contained in the **brainstem**. Among the BG structures, the most relevant to be studied at early developmental stages are the main input and output structures and their related cortical projections, implicating in particular the dorsal **striatum** that can be considered as the main input structure for SM projections, and the internal segment of the **globus pallidus** (GPi), one of the major output structures of the BG (Leisman et al., 2014).

Identifying these GM structures on MR images is quite challenging in infants due to the inter-subject variability and rapid changes in morphological characteristics and sizes during the perinatal period. To segment the **primary SM cortices**, the central sulcus was required as a landmark to delineate pre- and post- central regions; that is why we used a surfacic parcellation tool optimized for infants (MCRIB-S) (Adamson et al., 2020). After projecting these cortical ROIs to the cortical ribbon, a dilation into the WM was used to ease the tractography. For **sub-cortical structures**, a volumetric parcellation was used based on Draw-EM method (Makropoulos et al., 2014). It is worth noting that, even if the parcellation of the **subthalamic nucleus** was available, we decided not to use it in our study because its small size prevented an accurate fibers reconstruction. Also, we decided not to consider the **cerebellum**, whose undeniable yet little explored role in SM function during early development deserves a separate study.

#### 2. Reconstructing SM tracts of interest in infants

Individual dissections of SM connections, which to our knowledge have never been achieved in neonates and infants until now, were performed using an automated tractography-based approach benefitting from multi-shell MRI data. We optimized the settings (e.g., tractography parameters, exclusion mask) to reduce the number of spurious streamlines. We focused on a set of reliably delineated homotopic inter-hemispheric cortico-cortical and intra-hemispheric long-range cortico-subcortical tracts. Additionally, aware of the importance of the S1-M1 connectivity, intra-hemispheric short-range connections were also considered between pre- and post-central regions. Connections between subcortical structures were not studied because of their proximity which could alter the tractography performance. We visually validated the accuracy of the tracts reconstructions for several subjects and observed expected topographies (e.g., the S1 and M1 projections towards the ventral anterior and lateral portions of the thalami). We excluded the cortico-cortical tracts involving ParaC regions from the analyses due to the presence of frequent and variable tractography errors likely linked to their medial position. Overall, despite these limitations, we were confident in the quality of the reconstructed tracts.

#### 3. Evaluating the developing microstructure of SM tracts

We then quantified the microstructure of each SM tract by extracting DTI and NODDI-derived diffusion metrics using a weighting approach which privileges voxels with higher density. DTI and NODDI models present different trade-offs between complexity, biological plausibility, robustness, and run-time duration (Jelescu and Budde, 2017). Despite its role as a ‘ default’ choice in most studies in the context of WM development (Dubois et al., 2014; Ouyang et al., 2019), the DTI-derived metrics can be affected by several microstructural features and lack specificity to disentangle the complex properties of voxels containing crossing, kissing, and fanning fibers (Jeurissen et al., 2013; Zhang et al., 2012). On the reverse, NODDI allows a more sophisticated and biologically plausible multi-compartment model, relevant for developmental studies (Chang et al., 2015; Genc et al., 2017; Mah et al., 2017; Kimpton et al., 2021), but it requires multi-shell data and increased run-time duration, except when using GPU accelerations. In this study, we had to face some limitations related to the application of NODDI model to infant data. In particular, although potentially sub- optimal, we opted for default settings of diffusivities, which were optimized for the adult WM but not for infants (Guerrero et al., 2019). In the absence of gold standards for infant-specific NODDI fitting to evaluate the metric maps, this was performed to maintain some consistency with previous studies (Guerrero et al., 2019; Fenchel et al., 2020).

Overall, the resulting metrics maps appeared consistent across PT and FT subjects, and the metric differences across tracts seemed coherent between DTI and NODDI results in all infants (e.g., each cortico-brainstem tracts presenting a specific microstructural profile for all metrics but ODI). Globally, MD presented an opposite pattern to NDI (Kimpton et al., 2021) and RD seemed highly similar to MD in all tracts and opposite to FA in cortico-subcortical tracts, confirming that differences in MD and FA might be largely driven by RD. Such differences across tracts might result from differences in both intrinsic microstructure (similar to adults) and in maturation (according to different myelination stages across tracts) (Dubois et al., 2014). During WM development, MD tends to decrease with the growth of barriers to the random water motion, while FA tends to increase, reflecting several factors including the presence of compact fiber tracts and increasing myelination (Beaulieu and Allen, 1994). Additionally, previous studies suggested the accuracy of NODDI to assess WM maturation, with NDI being quite sensitive to WM microstructural developmental changes (Batalle et al., 2017; Kimpton et al., 2021). Besides, ODI describes the orientational dispersion of fibers within a tract, which is highly variable across tracts and might change during the maturation of crossing fibers (Raghavan et al., 2021). Nevertheless, additional analyses (not in the scope of this article) might be performed to evaluate the potential correlations between all these metrics, as performed in previous studies (Kunz et al., 2014).

Moreover, results on maturational distances (see II.4 for the discussion of the tracts comparisons) highlighted a higher coherence between DTI sets than with the NODDI set. In line with previous studies (Li et al., 2022), MD and FA did not bring additional benefits to the multimeric evaluations: consistency of tract differences evaluated by the two DTI sets confirmed the redundancy of information between the metric pairs. Additionally, the differences observed between DTI and NODDI sets (with NODDI set presenting more compact and roughly lower Mahalanobis distances values across tracts, with subtle variations in the main order) suggested that NODDI metrics provide complementary information for assessing microstructural maturational differences, probably due to their differing sensitivity of neurites to growth and maturation. Thus, confirming previous studies (Batalle et al., 2017; Kimpton et al., 2021), our results highlighted the complementarity of these models and confirmed the relevance of NODDI-derived metrics for the study of WM microstructure maturation in the context of prematurity.

### II. Studying the effects of prematurity on SM network maturation

#### 1. Study design

In contrast to most previous studies, we here focused on low-risk PT infants without overt brain abnormality of possible clinical significance, a cohort representative of the majority of children born prematurely nowadays in developed countries. Among the quite large FT cohort available in the dHCP, we considered the neonates with optimal pairing to PT infants, based on sex and age at MRI. Despite the presence of certain clinical risk factors for some PT infants (e.g., morbidities related to prematurity), the absence of significant difference between the BSID-III outcome of PT and FT infants at around 18 months corroborated that the included PT infants were at low-risk for neurodevelopment impairment. Furthermore, analyses on SM tracts were performed at TEA, age at which preterm infants are usually imaged with MRI so that they can be compared with typically developing FT neonates.

In this study, we hypothesized that the SM tracts would show a significant maturational delay in PT compared to FT neonates, with distinct patterns as a function of GA at birth and across cortico-subcortical and cortico-cortical tracts. This was based on the assumption that early peripheral stimuli are essential for the emerging SM network maturation, and that premature birth is associated with modified SM stimuli and experiences (notably related to numerous and various procedures in NICU (Mörelius et al., 2006; Gibbins et al., 2008), which might have a differential effect on somatosensory and motor systems (Duerden et al., 2018; Schneider et al., 2018; Jones et al., 2022). Of note, previous studies have reported either higher tactile sensitivity in PT infants at TEA (André et al., 2020), tactile hyporeactivity and/or undifferentiated integration of nociceptive versus non-nociceptive stimuli (Fabrizi et al., 2011), in association with some WM abnormalities (Brummelte et al., 2012; Zwicker et al., 2013).

#### 2. Classical analyses on the effects of prematurity

Univariate analyses were used to evaluate the effects of several factors on diffusion metrics measured in SM tracts at TEA. Although coherent with previous studies (Dubois et al., 2009; Liu et al., 2010; van Pul et al., 2012), inter-hemispheric asymmetries were not analyzed in details because this was out of the scope of this study, and the effects seemed similar across PT and FT infants. As expected, the metrics –even measured close to TEA– were dependent on PMA at MRI (de Bruïne et al., 2011; van Pul et al., 2012; Kimpton et al., 2021).

All SM tracts metrics except ODI were also related to the interindividual variability in whole-brain WM microstructure (beyond PMA at MRI), justifying the need to take these variables into account to highlight potential subtle effects.

The infant **group** had a significant influence on the metrics, with PT_EV_ infants showing more immature microstructural profiles (higher AD, RD, MD, lower FA, NDI, ODI) than PT_ML_ and FT infants. Interestingly, when analyses were performed at the tract level, PT_ML_ showed no difference with the FT paired group, suggesting that the specific SM tracts studied here might not contribute significantly to the well-described whole-brain WM diffusion abnormalities in moderate-late PT (Kelly et al., 2016a; Thompson et al., 2019). Once controlled for the effects of PMA at scan and WM microstructure, we observed that AD, RD, MD decreased with **GA at birth** (considered as a continuous variable) in all tracts, while FA, NDI and ODI increased. This was consistent with previous studies showing the same tendencies in most WM tracts with DTI and NODDI metrics, and suggesting more “immature” microstructural characteristics with higher prematurity degree (Kunz et al., 2014; Kelly et al., 2016a; Batalle et al., 2017; Thompson et al., 2019; Kimpton et al., 2021; Dibble et al., 2021). Interestingly, for a few tracts, we observed non-significant effects for FA and ODI, suggesting that both metrics might be less sensitive to detect variations of microstructure with GA at birth in the settings of this study.

Moreover, we explored the effects of additional factors on the metrics in SM tracts. Firstly, these were not related to the infants’ sex in our cohort. Combined with previous studies which showed inconsistent results (Pannek et al., 2014; Barnett et al., 2018; Kimpton et al., 2021), this observation suggests that sex effect might vary according to the studied tracts. Surprisingly, we observed no effect of main perinatal clinical risk factors, including preterm morbidities, despite numerous studies describing associations with WM abnormalities, notably for obstetric, neonatal and postnatal factors (Pogribna et al., 2013; Brouwer et al., 2017; Barnett et al., 2018; Parikh et al., 2021), and for exposure to cumulative risk factors (Barnett et al., 2018). Our negative results might be partly due to the study of low-risk PT infants.

As mentioned in the previous section, the metrics depended strongly on the tracts because of different microstructural and maturational properties. To evaluate the maturational gap related to prematurity at TEA, we visualized the relative percent changes of metrics in PT infants compared to paired FT neonates, but these remained difficult to interpret. We thus opted for an original multivariate approach to further compare PT and FT infants.

#### 3. Evaluating a maturational distance related to prematurity

Although univariate approaches might allow to make inferences on the effects of prematurity on the SM tract microstructure, no efficient and easily maturational interpretation can be inferred from single metrics which are sensitive to different underlying microstructural properties and maturational processes. To overcome this difficulty, we implemented an original approach, already validated in infants’ cohorts (Kulikova et al., 2015; Li et al., 2022) that took advantage of the complementary information described by different DTI and NODDI metrics, to enable better characterization of the SM tract maturation and inform on the effects of the prematurity as compared to typical development. Multivariate Mahalanobis distance was calculated in respect to a reference group (FT neonates) which provided typical values for the given tract. Importantly, this approach allows to consider inherent variability of the diffusion metrics across tracts in the FT group, and the correlations between the metrics, increasing the relevance of computed distances in the context of typical SM tract differences at TEA. To provide an accurate comparison of the metrics and groups, we scaled the metrics to the same range and corrected both PT and FT group values for the effects of GA at birth, PMA at scan, and whole-brain WM microstructure. For each tract, the resulting maturational distance related to prematurity could then be interpreted as a developmental gap between a PT infant at TEA and the FT control group.

#### 4. Highlighting the tract-specific effects of prematurity on SM network

Focusing on the PT_EV_ group, the comparison of distances across tracts highlighted the *differential impact of the prematurity on the SM tracts at TEA*. Whatever the set of diffusion metrics (see section I.3), the impact seems to increase in a **caudo-rostral and central-to-peripheral manner**, from the *cortico-Brainstem* tracts (having the smaller distances), followed by the tracts involving the *BG* and *thalamus* (with a greater distance specifically for the *Lenticular* tracts), then the *homotopic inter-hemispheric* tracts, and finally the *intra-hemispheric cortico-cortical S1-M1* tracts. Overall, this vulnerability pattern followed the typical progression of WM growth and myelination during infancy (Yakovlev and Lecours, 1967; Dubois et al., 2014) and within the CST tract (Kimpton et al., 2021), suggesting that tracts maturing early on are less impacted. Furthermore, while this spatial pattern is globally in line with previous studies on preterms (Wu et al., 2017; Knight et al., 2018), our results raised several interesting points regarding the functional role of the different SM tracts and related GM structures during development.

Firstly, we observed differences in maturational distances in tracts related to different cortical regions. **S1- subcortical** tracts seemed systematically more impacted by prematurity than M1- and ParaC-subcortical tracts. This suggests that S1 tracts may present a *specific vulnerability* to prematurity deleterious effects at TEA, in relation to altered SM perceptions and experiences. On the reverse, the observed differences may reflect a similar impact but a higher compensation of M1/ParaC-subcortical tracts following prematurity, relying on faster and/or more efficient maturational ‘ catch-up’ mechanisms during the first post-natal weeks.

The similar profiles of **ParaC-subcortical** tracts and related M1- tracts are less straightforward to interpret as the paracentral lobule includes both motor and somatosensory regions. Given the somatotopic organization of S1 and M1, this might suggest that connections related to regions of representation of lower limbs are less impacted by prematurity. Nevertheless, a methodological bias (more motor than sensory fibers included in ParaC- tracts) cannot be excluded.

Secondly, differences in maturational distances related to prematurity were also observed between tracts related to the different sub-cortical structures. The **cortico-Brainstem** tracts seemed the least impacted. Since it mainly consists of CST fibers that myelinate early on notably at the level of the PLIC (Dubois et al., 2014; Kulikova et al., 2015; Kimpton et al., 2021), this is coherent with the fact that connections with more advanced maturation at the time of birth might be less vulnerable to prematurity (Wu et al., 2017). The **cortico-Thalamic** tracts showed “intermediate” maturational distances in PT infants in reference to FT, which is in agreement with the acknowledged vulnerability of thalamocortical connectivity following preterm birth (Ball et al., 2013a). However, the specific functional position of the thalamus, with essential input and output projections to the different SM regions, may participate in modulating this vulnerability compared to other sub-cortical structures in relation to the early maturation of sensory functions in preterm infants (Duerden et al., 2018; Schneider et al., 2018). The **Cortico-Caudate** tracts showed similar “intermediate” maturational distances. This might result from an interplay between the high vulnerability to prematurity of caudate nuclei (Nosarti et al., 2014; Back, 2015; Loh et al., 2017), the alterations of the major efferent projections (McClendon et al., 2014) compared with more “preserved” afferences (from SM cortices) in relation to constant neonatal SM stimuli and experiences. Besides, the **Cortico-Lenticular** tracts systematically presented the greatest maturational distance, suggesting their particular vulnerability to prematurity. In addition to the known structural consequences of preterm birth on BG (Loh et al., 2017, 2020), different hypotheses can be proposed to explain this specific profile, especially knowing the anatomo-functional particularities of these tracts. As the dissected tracts is supposed to include *cortico-putaminal* (afferent) but also some (efferent) *pallido-cortical* fibers, the observed alteration may involve both input (putamen) and output (GPi) structures of the BG, which have different functions in cortico-BG loops. We may hypothesize that the maturation of the *efferent* pallido-cortical fibers is specifically altered by prematurity, with functional effects on the information reaching SM cortices and then, secondarily, inducing an altered maturation of the descending cortico-striatal and cortico-pallidal fibers. Finally, we observed globally that **cortico-cortical** tracts had higher maturational distances related to prematurity than cortico-subcortical tracts, suggesting a particular vulnerability which may be related to their later growth and maturation (Kostović et al., 2019). The *inter*-hemispheric tracts presented lower distances than *intra*-hemispheric S1-M1 tracts, in line with the late and protracted maturation of such connections.

While interesting, all these results should be interpreted cautiously given the limitations of diffusion MRI and tractography, in relation to the image spatial resolution, the size of neonatal structures and the presence of crossing fibers notably at the level of the corona radiata. Finally, it would be interesting to further investigate whether the vulnerability of SM tracts to prematurity is stable over development or whether some “catch-up” is present for some tracts, either before or after TEA. Nevertheless, this would require the longitudinal evaluation of maturational distances.

### III. Relating the early microstructure of SM tracts with neurodevelopmental outcome

#### 1. Study design

In this study, we intended to relate the SM microstructural characteristics at TEA to outcome at 18mCA, corresponding to a strategic age to assess severe neurodevelopmental impairments leading to some diagnoses (e.g., cerebral palsy). In this specific low-risk preterm cohort, no substantial developmental delay or specific disability was expected, as confirmed by the results. Similarly, in the case of significant alterations, we expected neuromotor impairments affecting more fine than gross motor skills, explaining our choice to analyze separated *scaled scores* (results for composite scores were nevertheless highly consistent).

#### 2. Relating the tracts maturational distances and outcome

We hypothesized correlations between diffusion metrics and BSID-III scores based on previous studies showing that, even in the absence of overt brain lesion, neonatal microstructural WM measures are associated with outcome in toddlers and children (van Kooij et al., 2012; Duerden et al., 2015; Barnett et al., 2018; Girault et al., 2019; Pannek et al., 2020; Kelly et al., 2020; Parikh et al., 2021). In particular, reduced FA (especially in the PLIC) has been associated with delayed psychomotor development and motor disability at different ages (Skranes et al., 2007; Rose et al., 2007; De Bruïne et al., 2013; Groeschel et al., 2014; Kelly et al., 2016b), and NODDI metrics have been related with neurodevelopmental outcomes (Kelly et al., 2016b; Young et al., 2019).

Our results showed negative correlations between **Cognitive** and **Fine motor scaled scores** in PT_EV_ and maturational distances for a number of tracts in **NODDI set** only. This suggested that the early microstructural information coded by NODDI might be more relevant than DTI for detecting subtle tract alterations related to later neurodevelopmental impairments. This is in line with previous studies showing that NODDI is more sensitive than FA to disruptions in WM development in preterms (Batalle et al., 2017; Kimpton et al., 2021). Moreover, as hypothesized, early SM tract microstructure was further correlated with *cognitive* performances during toddlerhood, confirming the essential developmental interactions between the SM system and higher-order functions, and the common clinical overlap of motor and cognitive impairments in the PT population.

Regarding the specific tracts showing correlations with outcome in the PT_EV_ group, we first observed that **fine motor** score was related to 5 tracts presenting different profiles of maturational distances: ***M1-Brainstem and ParaC-Brainstem*** with the lowest distances; ***M1-Lenti and ParaC-Lenti*** with intermediate distances; and ***S1- M1*** with the greatest distances. This suggested that the degree of maturational gap at TEA by itself is not the only factor explaining the motor performances.

In the light of our results showing the high vulnerability of *lenticular* tracts to prematurity, it is not surprising that microstructural alterations in the motor tracts connected to this key BG structure may underpin early SM impairments with further consequences on fine motor skills acquisitions (Leisman et al., 2014). Likewise, as the intra-hemispheric SM connections contribute to improve SM integration and functions, the correlation observed for *S1-M1* tract suggests that the early prematurity impact on these tracts may alter neuromotor development with discernible impairments in toddlers.

Besides, distances for ***M1-Brainstem and ParaC-Brainstem*** tracts were also correlated with **Cognitive** scores, which is interesting knowing the functional importance of *brainstem* for SM and non-SM functions: it is then easy to imagine that the early microstructural alterations in the correspondent tracts can be associated with functional alterations with consequences on global neurodevelopmental acquisitions, including both cognitive and motor performances.

These relationships observed in PT_EV_ between SM tract microstructure at TEA and outcome at 18mCA are of particular interest in the prematurity context, as even low-risk populations are at increased risk of (sometimes subtle) neuromotor disorders (e.g., developmental coordination disorder) (Edwards et al., 2011; Spittle and Orton, 2014; Zwicker, 2014; Groeschel et al., 2019). However, these disorders are not visible enough to be diagnosed until much later (often at school age) (Williams et al., 2010; de Jong et al., 2012; Van Hus et al., 2014), which underlines the need for early biomarkers. Thus, the specific NODDI maturational profile of these 5 primary SM tracts should be further explored, in order to investigate their potential value as early markers of motor and/or other neurodevelopmental disorders such as developmental coordination disorder. Nevertheless, although relating early brain markers and long-term outcome has important clinical relevance, we should remind that, as children grow older, environmental factors (e.g., socio-familiar context) seem to explain the greatest part of interindividual variability in neurodevelopment, with the influence of perinatal risk factors diminishing over time (Thompson Jr. et al., 1998; Miceli et al., 2000; Anderson and Doyle, 2008; Linsell et al., 2015). Thus, future studies should incorporate more accurate predictive models to intend to approach the complex relationship between early brain characteristics and outcome.

## Conclusions

Using a combination of innovative methods, our results confirmed that prematurity impacts the early microstructural development of the primary SM network at TEA, even in low-risk preterm infants. We further highlighted those effects differ according to GA at birth, but also across the SM tracts, with the more rostral tracts as well as tracts involving S1 showing the greatest vulnerability to prematurity noxious effects. Our study also showed the complementarity between DTI and NODDI models as well as the interest of using multiparametric approaches for assessing maturational processes and microstructural developmental differences. Longitudinal studies including earlier MRI evaluations as well as behavioral follow-up until later ages would provide a better understanding of the impact of early-life disturbances in SM tracts microstructure on neurodevelopmental disorders.

## Abbreviations

AD: Axial diffusivity
BG: basal ganglia
BSID-III: Bayley Scales of Infant and toddler Development, 3^rd^ edition
Bstem: brainstem
Caud: caudate nucleus
CLD: chronic lung disease
CSF: cerebrospinal fluid
CST: corticospinal tract
cUS: cranial ultrasonography
dHCP: Developing Human Connectome Project
dMRI: Diffusion-weighted MRI
DTI: Diffusion tensor imaging
FA: Fractional anisotropy
FDR: False Discovery Rate
FT: full-term born infants
[**FT**_**EVCt**_: ‘ extreme to very preterms’ matched FT controls,
**FT**_**MLCt**_: ‘ moderate to late preterms’ matched FT controls]
GA: gestational age
GM: gray matter
IMD: Index of multiple deprivation
IUGR: intrauterine growth restriction
Lenti: lentiform nucleus
M1: lateral portion of the primary motor cortex (precentral region)
mCA: months of corrected age
MD: Mean diffusivity
MRI: magnetic resonance imaging
NDI: Neurite Density Index
NEC: Necrotizing enterocolitis
NICU: neonatal intensive care unit
NODDI: Neurite Orientation Dispersion and Density Imaging
ODI: Orientation Dispersion Index
ParaC: medial portions of the primary sensori-motor cortices (paracentral region)
PMA: post-menstrual age
PT: preterm infants
[**PT**_**EV**_: extreme to very preterms,
**PT**_**ML**_: moderate to late preterms]
RD: Radial diffusivity
ROIs: regions of interest
ROP: Retinopathy of prematurity
S1: lateral portion of the primary somatosensory cortex (postcentral region)
SES: Socio-Economic Status
Thal: thalamus
TEA: term- equivalent age
wGA: weeks of GA
WM: white matter

## Author Contributions

*Conceptualization*: SN, AG, LHP, TA and JD; *Methodology*: SN, AG, YL and JD; *Resources and Investigation*: SN, AG, TA and JD; *Data curation*: SN, AG; *Validation and Formal Analysis*: SN, AG and JD; *Software*: AG and YL; *Supervision*: TA and JD; *Visualization*: SN, AG and JD; *Writing – original draft*: SN, AG and JD; *Writing – review & editing*: SN, AG, YL, LHP, TA and JD.

## Acknowledgments

We thank Jean-François Mangin for the methodological support; Nicholas Harper for providing the individual clinical data; and the infants and their families for their participation in this study.

