## Supplementary Material for "Early structural connectivity within the sensorimotor network: deviations related to prematurity and association to neurodevelopmental outcome"

### Supplementary Tables

**Table SupT1. Clinical factors for the 4 infants' sub-groups used for the adjustments in descriptive models.** Preterm infants (PT) were subdivided into those born extremely to very preterm ($PT_{EV}$) or moderate to late preterm ($PT_{ML}$) and their corresponding full-term (FT) controls.

| | **$PT_{EV}$** (N=33) | **$FT_{EVCt}$** (N=33) | **$PT_{ML}$** (N=26) | **$FT_{MLCt}$** (N=26) |
|---|---|---|---|---|
| GA at birth (*weeks*) *Median (IQR) [range]* | 29.0 (27.4; 30.7) [23.7; 31.9] | 40.4 (39.9; 41.1) [37.4; 42.3] | 34.3 (33.2; 35.1) [32.3; 36.0] | 40.1 (39.4;40.8) [37.4;42.1] |
| Sex, *male* | 18 (55%) | 18 (55%) | 15 (58%) | 15 (58%) |
| Multiple Pregnancy, *twins* | 9 (27%) | 0 | 9 (35%) | 1 (4%) |
| IUGR | 10 (33%) *NA: 3* | 0 *NA: 1* | 7 (28%) *NA: 1* | 1 (4%) *NA: 1* |
| Preterm Morbidities | 21 (64%) | 0 | 4 (36%) *NA: 15* | 0 |
| Parenteral Nutrition >21days | 4 (12%) | 0 | 0 *NA: 5* | 0 |
| PMA at MRI (*weeks*) *Median (IQR) [range]* | 41.3 (40.1; 42.3) [38.4; 44.9] | 41.3 (40.3;42.3) [38.3;44.7] | 41.1 (40.1;42.0) [38.9;44.1] | 41.1 (40.1;42.0) [38.9;44.1] |

Refer to *Table 1* legend for abbreviations.

**Table SupT2. Cohort characteristics and outcome assessment at around 18 months of corrected age.**

| | | **Preterm infants** (*n*=59) | | **Full-term controls** (*n*=59) | | |
|---|---|---|---|---|---|---|
| | | *NA* | | *NA* | | *p* |
| **Sex**, male | N (%) | *15* | 23 (52.3) ¤ | *6* | 30 (56.6) ¤ | *#* |
| **Age at assessment** (*months*) | Median (IQR) [Range] | *15* | | *6* | | |
| *Chronological* age | | | 20.4 (19.8;21.1) [18.9;23.4] | | 18.2 (17.8;18.7) [17.3;19.8] | *** |
| *Corrected* age | | | 18.4 (18.1;18.7) [17.7;20.8] | | 18.2 (17.9;18.6) [17.3;19.8] | ns |
| **Family socioeconomic status:** Index of multiple deprivation (IMD) | Median (IQR) [Range] | *17* | 17.8 (11.1;29.7) [2.7;48.3] | *5* | 26.6 (18.2;33.8) [4.2;46.4] | * |
| BSID-III: ***Scaled* scores** | Mean (SD) [range] | *15* | | *6* | | |
| Cognitive | | | 10.1 (2.6) [1.0;15.0] | | 10.2 (2.0) [4.0;15.0] | ns |
| Receptive communication | | | 10.7 (3.8) [1.0;19.0] | | 10.8 (3.1) [5.0;18.0] | ns |
| Expressive communication | | | 8.7 (2.8) [1.0;15.0] | | 9.6 (2.5) [4.0;14.0] | ns |
| Fine motor | | | 11.0 (2.5) [2.0;15.0] | | 11.1 (2.2) [4.0;16.0] | ns |
| Gross motor | | | 9.0 (2.0) [3.0;16.0] | | 9.3 (1.6) [5.0;13.0] | ns |

¤ percentage over the available data (see NA for missing data). Comparisons for ages at assessment and IMD scores were performed with Wilcoxon rank sum test. #: no comparisons done for sex (used for pairing the full-term controls). *p*-values for BSID-III scores were obtained from t-tests corrected for multiple comparisons. Refer to *Table 1* legend for abbreviations. See *Figure 2* for BSID-III scores distributions and graphical comparison between groups.

**Tables SupT3. A.** ANOVA model studying effects of tract, hemisphere, infant group (considering *extreme to very preterms* ($PT_{EV}$) / *moderate to late preterms* ($PT_{ML}$) and their paired controls ($FT_{EVCt}$ and $FT_{MLCt}$, respectively), and their interactions on each diffusion metric. **B.** Paired t-test comparisons between $PT_{EV}$, $PT_{ML}$, and paired FT controls for each diffusion metric, over all tracts (p-values corrected for multiple comparisons).

Refer to *Table 1* for abbreviations and *p*-values significance.

**A.**

| | AD | RD | MD | FA | NDI | ODI |
|---|---|---|---|---|---|---|
| Tract | **** | **** | **** | **** | **** | **** |
| Hemisphere | ns | * | ns | *** | ns | **** |
| Group | **** | *** | **** | ns | *** | **** |
| Tract : Hemisphere | **** | * | **** | **** | **** | **** |
| Tract : Group | **** | **** | **** | **** | **** | **** |
| Hemisphere : Group | ns | ns | ns | ns | ns | ns |

**B.**

| | $PT_{EV}$ *vs* $FT_{EVCt}$ (N= 33) | | | $PT_{ML}$ *vs* $FT_{MLCt}$ (N = 26) | | |
|---|---|---|---|---|---|---|
| | T | CI95% | *p*-value | T | CI95% | *p*-value |
| **AD** | 22.49 | [0.; 0.] | **** | 6.02 | [0.;0.] | **** |
| **RD** | 20.98 | [0.; 0.] | **** | 5.49 | [0.;0.] | **** |
| **MD** | 22.73 | [0.; 0.] | **** | 5.87 | [0.;0.] | **** |
| **FA** | -11.47 | [-0.02;-0.01] | **** | -2.11 | [-0.01;0.] | ** |
| **NDI** | -21.76 | [-0.03;-0.02] | **** | -3.78 | [-0.01;0.] | *** |
| **ODI** | -13.84 | [-0.02;-0.01] | **** | -4.87 | [-0.01;0.] | **** |

**Table SupT4**. **Group comparisons per tract**: paired t-tests for each diffusion metric between $PT_{EV}$ vs $FT_{EVCt}$ (N=33 in each group). Left/Right tracts values are averaged, as for all the further supplementary tables. P-values are corrected for multiple comparisons.

| | AD | | RD | | MD | | FA | | NDI | | ODI | |
|---|---|---|---|---|---|---|---|---|---|---|---|---|
| **Tracts** | T | *p* | T | *p* | T | *p* | T | *p* | T | *p* | T | *p* |
| S1-Bstem | 5.39 | **** | 3.9 | *** | 5.01 | **** | -0.6 | ns | -3.28 | ** | -4.87 | **** |
| M1-Bstem | 5.0 | **** | 4.78 | **** | 5.42 | **** | -1.92 | . | -4.22 | *** | -3.09 | ** |
| ParaC-Bstem | 4.55 | *** | 6.11 | **** | 6.22 | **** | -3.53 | ** | -5.89 | **** | -1.83 | . |
| S1-Thal | 7.48 | **** | 5.6 | **** | 6.34 | **** | -2.6 | * | -6.22 | **** | -6.31 | **** |
| M1-Thal | 6.32 | **** | 4.7 | **** | 5.36 | **** | -1.88 | . | -5.34 | **** | -5.01 | **** |
| ParaC-Thal | 4.58 | **** | 4.94 | **** | 4.97 | **** | -3.55 | ** | -5.3 | **** | -2.91 | ** |
| S1-Caud | 8.12 | **** | 4.5 | *** | 5.76 | **** | -0.24 | ns | -5.42 | **** | -5.36 | **** |
| M1-Caud | 5.47 | **** | 2.93 | ** | 3.7 | ** | 0.08 | ns | -3.78 | *** | -6.16 | **** |
| ParaC-Caud | 5.23 | **** | 5.94 | **** | 6.05 | **** | -3.67 | ** | -6.35 | **** | -3.31 | ** |
| S1-Lenti | 6.84 | **** | 6.18 | **** | 6.52 | **** | -4.6 | **** | -6.15 | **** | -6.12 | **** |
| M1-Lenti | 5.62 | **** | 5.76 | **** | 5.91 | **** | -4.1 | *** | -5.98 | **** | -3.36 | ** |
| ParaC-Lenti | 6.11 | **** | 6.04 | **** | 6.33 | **** | -4.03 | *** | -6.47 | **** | -2.57 | * |
| $S1_L$-$S1_R$ | 5.75 | **** | 9.23 | **** | 9.08 | **** | -6.73 | **** | -7.86 | **** | -0.91 | ns |
| $M1_L$-$M1_R$ | 5.26 | **** | 8.23 | **** | 7.99 | **** | -6.39 | **** | -7.22 | **** | -3.64 | ** |
| S1-M1 | 6.94 | **** | 7.42 | **** | 7.4 | **** | -5.49 | **** | -7.24 | **** | -3.64 | ** |

Positive T values refer to higher values in FT than PT. Refer to *Table 1* for *p*-value legend and to *Figure 1* for metrics and tracts abbreviations.

**Table SupT5.** ANOVA analysis including only the preterm infants ($PT_{EV}$ + $PT_{ML}$) to study the effect of selected clinical factors on each diffusion metric.

| | AD | RD | MD | FA | NDI | ODI |
|---|---|---|---|---|---|---|
| Tract | **** | **** | **** | **** | **** | **** |
| Group | * | ns | . | ns | . | * |
| Sex | ns | ns | ns | ns | ns | ns |
| Multiple Pregnancy | ns | ns | ns | ns | ns | * |
| IUGR | * | . | * | ns | * | * |
| Preterm Morbidities | ns | ns | ns | ns | ns | ns |
| Parenteral Nutrition >21d | * | . | . | ns | * | * |
| Tract : Group | ns | ** | * | **** | ns | ns |
| Group : Sex | ns | ns | ns | ns | ns | ns |
| Group : Multiple Pregnancy | ns | ns | ns | ns | ns | * |
| Group : IUGR | ns | ns | ns | ns | ns | * |
| Group : Preterm Morbidities | ns | ns | ns | ns | ns | ns |
| Group : Parenteral Nutrition | NA | NA | NA | NA | NA | NA |
| Sex : Multiple Pregnancy | ns | ns | ns | ns | ns | ns |
| Sex : IUGR | ns | ns | ns | ns | ns | ns |
| Sex : Preterm Morbidities | ns | ns | ns | ns | ns | ns |
| Sex : Parenteral Nutrition | ns | ns | ns | ns | ns | ns |

Refer to *Table 1* legend for abbreviations and *p*-values significance.

**Tables SupT6. A.** ANCOVA analysis including GA at birth (rather than the group belonging), PMA at scan, WM residuals (corrected for GA at birth and PMA at scan), relevant clinical factors (sex, multiple pregnancy, IUGR, parenteral nutrition >21days), as well as relevant interactions between factors. **B.** Multiple linear regression over the whole cohort of mean WM metrics with PMA at scan and GA at birth, and interactions.

Refer to *Table 1* legend for abbreviations and *p*-values significance.

**A.**

| | AD | RD | MD | FA | NDI | ODI |
|---|---|---|---|---|---|---|
| Tract | **** | **** | **** | **** | **** | **** |
| GA at birth | **** | **** | **** | **** | **** | **** |
| PMA at scan | **** | **** | **** | **** | **** | * |
| Residual WM | **** | **** | **** | **** | **** | **** |
| Sex | ns | ns | ns | ns | ns | ns |
| Multiple Pregnancy | ns | ns | ns | ns | ns | ns |
| IUGR | * | * | * | . | . | ns |
| Parenteral Nutrition | ns | ns | ns | ns | ns | ns |
| Tract : GA at birth | **** | **** | **** | **** | **** | **** |
| Tract : PMA at scan | *** | **** | **** | **** | **** | ns |
| Tract : Residual WM | **** | **** | **** | **** | **** | ns |
| Tract : Sex | * | ns | ns | ns | ns | * |
| Tract : Multiple Pregnancy | ns | ** | ns | **** | ns | ns |
| Tract : IUGR | ns | ns | ns | ns | ns | ns |
| Tract : Parenteral Nutrition | ns | **** | ** | ** | ** | ns |

**B.**

| | AD | RD | MD | FA | NDI | ODI |
|---|---|---|---|---|---|---|
| PMA at scan | **** | **** | **** | **** | **** | **** |
| GA at birth | **** | **** | **** | **** | **** | * |
| PMA : GA at birth | ns | ns | ns | ns | ns | ns |

**Table SupT7.** ANOVA model studying effects of tract and PT subgroup on Mahalanobis distances, for each set.

| | Set 1 (AD, RD) | Set 2 (MD, FA) | Set 3 (NDI, ODI) |
|---|---|---|---|
| Tract | **** | **** | **** |
| Group | **** | **** | **** |
| Tract : Group | **** | **** | **** |

Refer to *Table 1* for *p*-value legend.

**Table SupT8. Pearson correlation analyses between Mahalanobis distance and BSID-III scaled scores .** Only $PT_{EV}$ results for *set 3* (NODDI) are shown (p-values corrected for multiple comparisons), as no significant correlations were found in *sets 1* and *2*.

| | **$PT_{EV}$ - Set 3 (NDI, ODI)** | | | | | | | | | |
|---|---|---|---|---|---|---|---|---|---|---|
| | **Cognitive** | | **Receptive Com.** | | **Expressive Com.** | | **Fine Motor** | | **Gross Motor** | |
| Tract | r | *p* | r | *p* | r | *p* | r | *p* | r | *p* |
| S1-Bstem | -0.51 | ns | -0.46 | ns | -0.31 | ns | -0.56 | . | 0.12 | ns |
| **M1-Bstem** | -0.7 | * | -0.55 | . | -0.5 | ns | -0.68 | * | 0.08 | ns |
| **ParaC-Bstem** | -0.66 | * | -0.45 | ns | -0.47 | ns | -0.69 | * | 0.03 | ns |
| S1-Thal | -0.35 | ns | -0.22 | ns | -0.13 | ns | -0.48 | ns | 0.33 | ns |
| M1-Thal | -0.5 | ns | -0.43 | ns | -0.31 | ns | -0.58 | . | 0.08 | ns |
| ParaC-Thal | -0.34 | ns | -0.32 | ns | -0.25 | ns | -0.54 | ns | 0.1 | ns |
| S1-Caud | -0.41 | ns | -0.12 | ns | 0.02 | ns | -0.31 | ns | 0.13 | ns |
| M1-Caud | -0.27 | ns | -0.15 | ns | -0.12 | ns | -0.33 | ns | 0.12 | ns |
| ParaC-Caud | -0.35 | ns | -0.39 | ns | -0.33 | ns | -0.42 | ns | -0.02 | ns |
| S1-Lenti | -0.52 | ns | -0.48 | ns | -0.36 | ns | -0.57 | . | 0.15 | ns |
| **M1-Lenti** | -0.46 | ns | -0.51 | ns | -0.4 | ns | -0.62 | * | 0.02 | ns |
| **ParaC-Lenti** | -0.45 | ns | -0.47 | ns | -0.41 | ns | -0.62 | * | -0.02 | ns |
| $S1_L$-$S1_R$ | -0.01 | ns | -0.02 | ns | 0.11 | ns | -0.06 | ns | 0.17 | ns |
| $M1_L$-$M1_R$ | -0.47 | ns | -0.49 | ns | -0.43 | ns | -0.56 | . | -0.03 | ns |
| **S1-M1** | -0.56 | . | -0.54 | ns | -0.35 | ns | -0.68 | * | 0.1 | ns |

Refer to *Table 1* for p-value legend and to *Figure 1* for tracts abbreviations.

#### Supplementary Figures

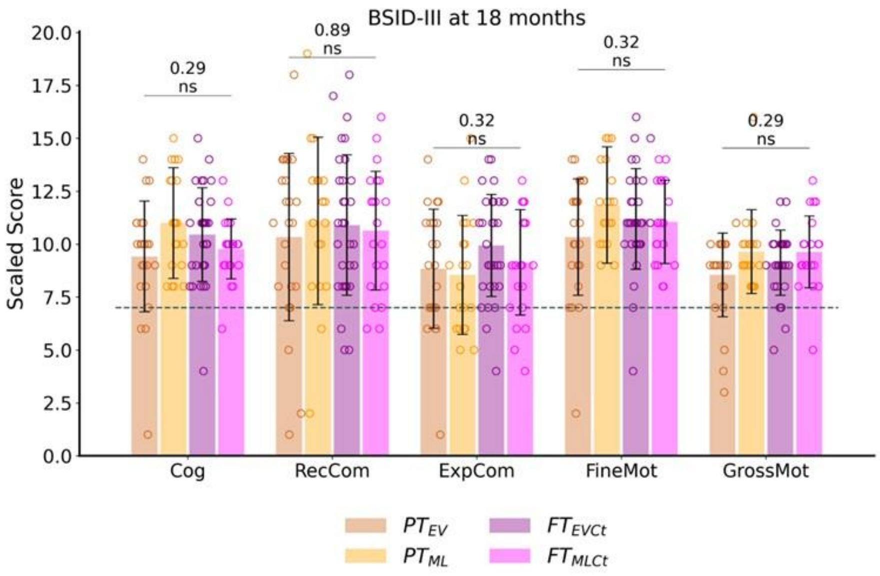


**Figure SupF1**. **Outcome assessment at around 18 months of corrected age: BSID-III scaled scores across subgroups** ($PT_{EV}$, $PT_{ML}$, $FT_{EVCt}$, and $FT_{MLCt}$), with the results of one-way ANOVA and *p*-values indicating no significant group effect. The dotted line indicates the scores threshold indicating a developmental delay (scores < -1 SD): scaled scores < 7. Cog: cognitive; RecCom: receptive communication, ExpCom: expressive communication; FineMot: fine motor, GrossMot: gross motor scores. ns: not significant.

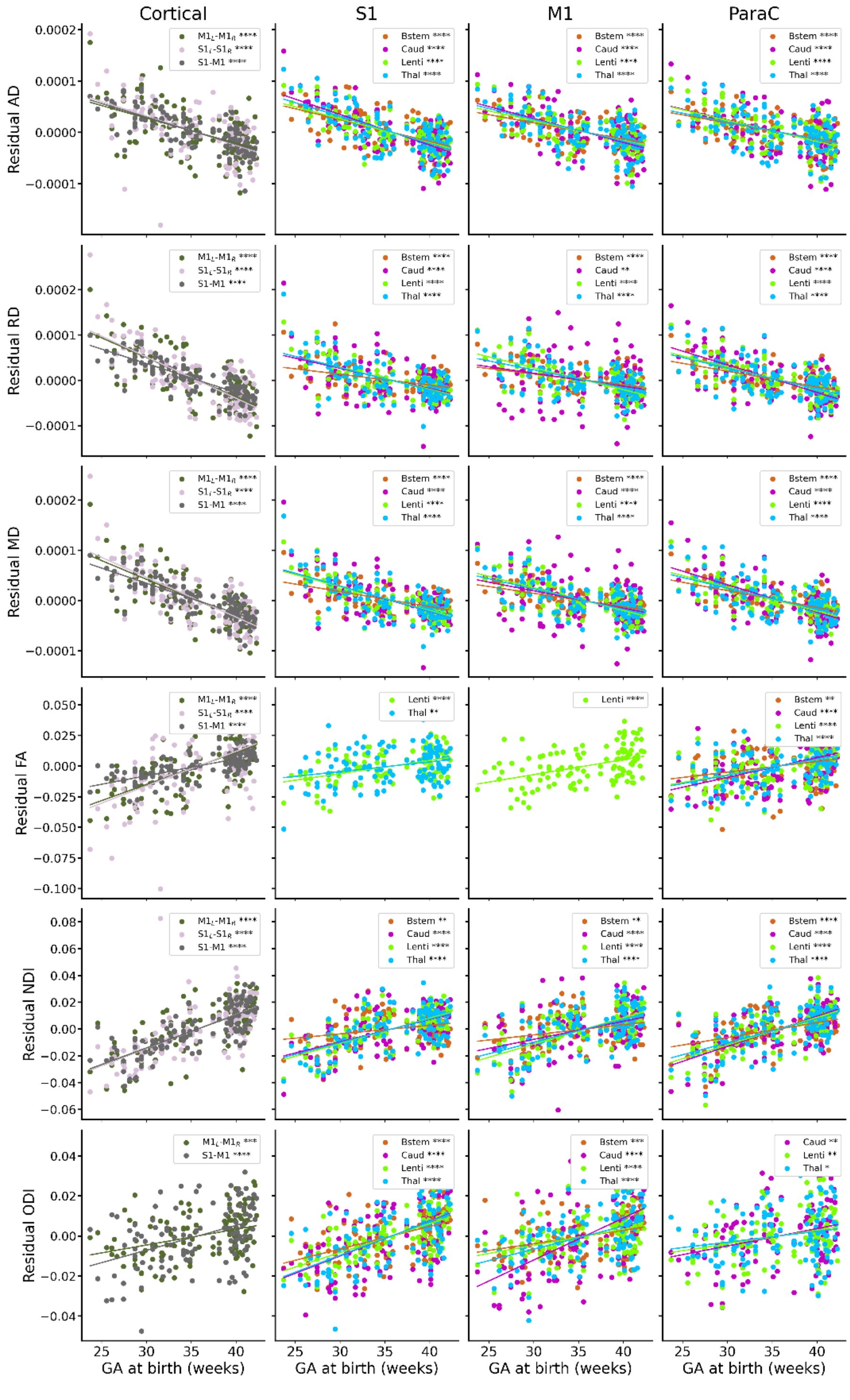


| *Metric* | *Tracts not significantly associated with GA at birth* |
|---|---|
| AD | - |
| RD | - |
| MD | - |
| FA | S1-Bstem<br>M1-Bstem<br>M1-Thal<br>S1-Caud<br>M1-Caud |
| NDI | - |
| ODI | ParaC-Bstem<br>$S1_L$-$S1_R$ |

**Figure SupF2. Scatterplot of metrics residuals (after correction of PMA at scan and WM residuals) with GA at birth**. The regression lines show significant relationship with GA at birth (after correction for multiple comparisons). The table lists the tracts that are not significantly associated with GA at birth for each metric. Refer to *Figure 1* legend for abbreviations.

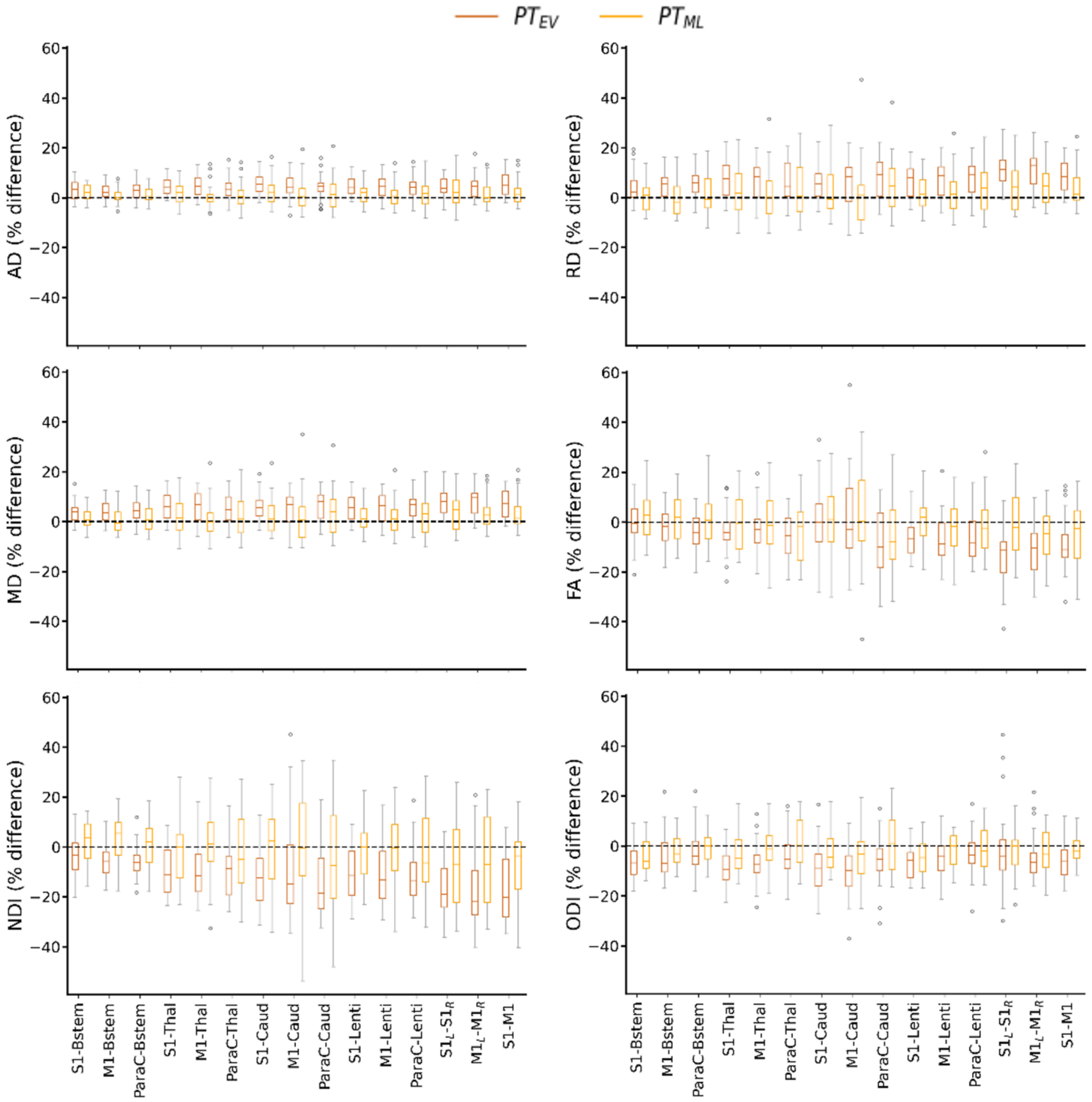


**Figure SupF3.** Relative percent difference in diffusion metrics between $PT_{EV}$ vs $FT_{EVCt}$ and $PT_{ML}$ vs $FT_{MLCt}$ groups (each PT infant being compared to his/her paired FT newborn) for each SM tract. Refer to *Figure 1* legend for abbreviations.

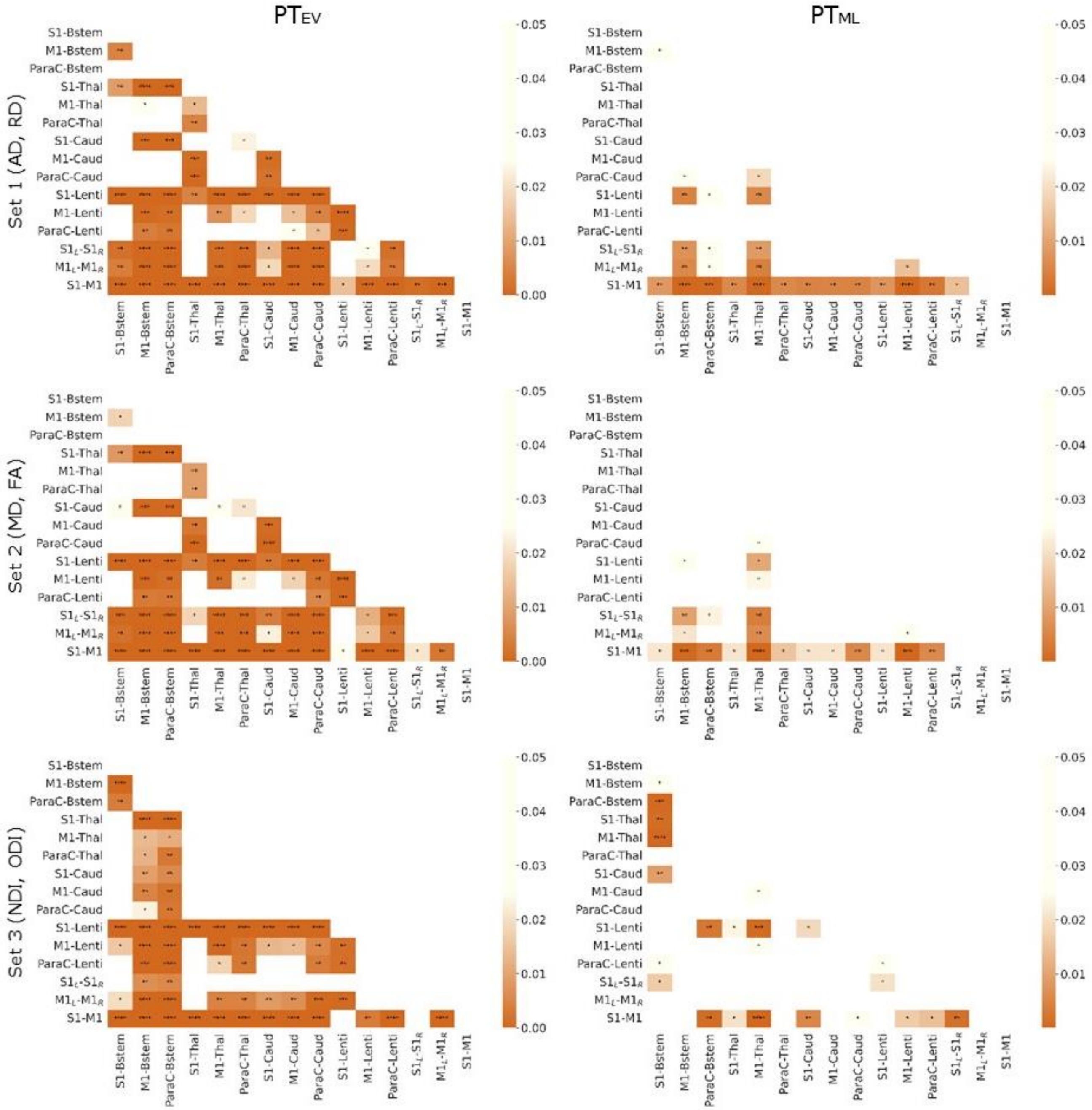


**Figure SupF4. Differential effect of prematurity on specific SM tracts.** Heatmap of p-values from paired t-tests (corrected for multiple comparisons) of Mahalanobis distances for each pair of SM tracts, for each set of metrics, in each PT subgroup independently ($PT_{EV}$ on the left, $PT_{ML}$ on the right). Legend depicts *p*-values with respect to heat intensity, with darker colours representing greater statistical significance and white no significant differences. Refer to *Figure 1* legend for abbreviations and to *Table 1* for p-value legend.

### Supplementary Materials

#### *Results for BSID-III composite score (CS)*

Composite scores below 85 (corresponding to <-1SD, indicating a developmental delay) concerned 16.5% of infants for Communication (N=16, 9 PT), 6.2% for Cognition (N=6, 4 PT) and 5.2% for Motricity (N=5, 3 PT).

**Table Sup-CS_T1**. Comparison of BSID-III composite scores at around 18 months of corrected age between PT and FT infants. p-values were obtained from paired t-tests corrected for multiple comparisons. See *Figure Sup-CS-F1* for distributions and graphical comparison between groups.

| | **Preterm group** (*n*=59) | | **Full-term controls** (*n*=59) | | |
|---|---|---|---|---|---|
| **Bayley's scores** (BSID-III) | *NA* | Mean (SD) [range] | *NA* | Mean (SD) [range] | *p* |
| ***Composite* scores** | *15* | | *6* | | |
| Cognitive | | 100.7 (13.1) [55.0;125.0] | | 100.8 (9.8) [70.0;125.0] | ns |
| Communication | | 98.4 (18.3) [47.0;141.0] | | 101.4 (15.5) [68.0;127.0] | ns |
| Motor | | 100.4 (10.5) [70.0;124.0] | | 101.6 (8.5) [73.0;118.0] | ns |

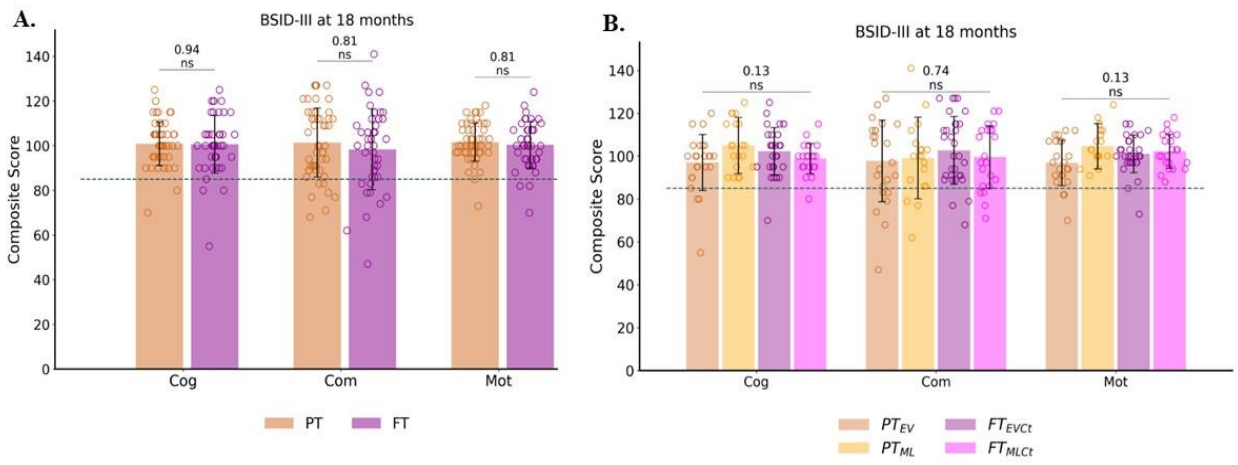


**Figure Sup-CS_F1.** Outcome assessment at around 18 months of corrected age: BSID-III composite scores between PT and FT groups (**A**), and subgroups (**B**: $PT_{EV}$, $PT_{ML}$, $FT_{EVCt}$ and $FT_{MLCt}$).

The dotted line indicates the scores threshold indicating a developmental delay (scores < -1 SD): composite scores < 85. Reported statistics are results of either t-test corrected for multiple comparisons for PT vs FT or one-way ANOVA for the subgroup analysis. Cog: cognitive; Com: communication; Mot: motor. ns: not significant.

**Table Sup-CS_T2.** Correlation analyses between Mahalanobis distance and BSID-III composite scores**.** Only $PT_{EV}$ results for *set 3* (NDI and ODI) are shown (p-values corrected for multiple comparisons), as no significant correlations were found in *sets 1* and *2*.

| | **$PT_{EV}$ - Set 3 (NDI, ODI)** | | | | | |
|---|---|---|---|---|---|---|
| | **Cognitive** | | **Communication** | | **Motor** | |
| **Tract** | r | *p* | r | *p* | r | *p* |
| S1-Brainstem | -0.51 | ns | -0.41 | ns | -0.37 | ns |
| **M1-Brainstem** | -0.7 | * | -0.55 | ns | -0.49 | ns |
| **ParaC-Brainstem** | -0.66 | * | -0.48 | ns | -0.52 | ns |
| S1-Thal | -0.35 | ns | -0.19 | ns | -0.18 | ns |
| M1-Thal | -0.5 | ns | -0.4 | ns | -0.4 | ns |
| ParaC-Thal | -0.34 | ns | -0.3 | ns | -0.35 | ns |
| S1-Caud | -0.41 | ns | -0.06 | ns | -0.16 | ns |
| M1-Caud | -0.27 | ns | -0.14 | ns | -0.18 | ns |
| ParacC-Caud | -0.35 | ns | -0.38 | ns | -0.33 | ns |
| S1-Lenti | -0.52 | ns | -0.45 | ns | -0.36 | ns |
| M1-Lenti | -0.46 | ns | -0.49 | ns | -0.47 | ns |
| ParaC-Lenti | -0.45 | ns | -0.47 | ns | -0.49 | ns |
| S1L-S1R | -0.01 | ns | 0.03 | ns | 0.04 | ns |
| M1L-M1R | -0.47 | ns | -0.49 | ns | -0.45 | ns |
| S1-M1 | -0.56 | ns | -0.47 | ns | -0.48 | ns |

Refer to *Table 1* for p-value legend and to *Figure 1* for tracts abbreviations.
